# “Abnormal vertebral patterns in genetically heterogeneous deceased fetuses and neonates: evidence of selection against variations”

**DOI:** 10.1101/784926

**Authors:** Pauline C. Schut, Erwin Brosens, Frietson Galis, Clara M. A. Ten Broek, Inge M.M. Baijens, Marjolein H.G. Dremmen, Dick Tibboel, Martin P. Schol, Annelies De Klein, Alex J. Eggink, Titia E. Cohen-Overbeek

## Abstract

**Objective:** To assess the vertebral pattern in a cohort of deceased fetuses and neonates, and to study the possible impact of DNA Copy Number Variations (CNVs) in coding regions and/or disturbing enhancers on the development of the vertebral pattern.

**Method:** Radiographs of 445 fetuses and infants, deceased between 2009 and 2015, were assessed. Terminations of pregnancies, stillbirths and neonatal deaths were included. Patients were excluded if the vertebral pattern could not be determined. Copy number profiles of 265 patients were determined using single nucleotide polymorphism array.

**Results:** 274/374 patients (73.3%) had an abnormal vertebral pattern. Cervical ribs were present in 188/374 (50.3%) and were significantly more common in stillbirths (69/128 (53.9%)) and terminations of pregnancies (101/188 (53.7%)), compared to live births (18/58, 31.0%, p = 0.006). None of the rare CNVs were recurrent or overlapped candidate genes for vertebral patterning.

**Conclusion:** The presence of an abnormal vertebral pattern, particularly in the cervical region, could be a sign of disruption at critical, highly interactive and conserved stages of embryogenesis. The vertebral pattern might provide valuable information regarding fetal and neonatal outcome. CNV analyses did not identify a mutual genetic cause for the occurrence of vertebral patterning abnormalities, indicating genetic heterogeneity.

## Introduction

The human vertebral column normally consists of 7 cervical, 12 thoracic, 5 lumbar, 5 sacral and 3-4 coccygeal vertebrae. Only the thoracic vertebrae are rib-bearing. Deviations from this vertebral pattern rarely occur in healthy individuals, particularly in the cervical region.^1^ However, variations in cervical patterning, including (rudimentary) cervical ribs, have been described in specific populations.^2,3^ In the presence of cervical ribs, a partial or full posterior homeotic transformation of the seventh cervical vertebra has occurred, because the vertebra has features of a rib-bearing thoracic vertebra. This results in a change in the number of true cervical and thoracic vertebrae and consequently to a shift of the cervicothoracic junction.^4^

It has been hypothesized that the lack of variation in cervical vertebral patterning is the result of developmental constraints or evolutionary selection against changes.^5-8^ The low prevalence of cervical ribs in healthy pediatric or adult populations, compared to the high prevalence in deceased fetuses and neonates, supports this hypothesis.^7,9^ A high prevalence of abnormalities in vertebral patterning is also found in children with specific pediatric malignancies.^10-13^ Furthermore, Galis *et al.* ^7^ concluded that the majority of individuals with cervical ribs, are not expected to reach reproductive age. While the presence of cervical ribs is usually not directly life-threatening, cervical ribs could be regarded as markers of disadvantageous embryonic development, which in turn can result in an adverse outcome.^14^ The underlying causal mechanism of abnormalities in vertebral patterning and adverse developmental effects is currently unknown, but the strong interactions between anterior-posterior patterning and the development of different organ systems at early embryonic stages could play a role.^6^ Disruption of the vertebral pattern at the cervical level is expected to be more harmful than disruptions at thoracic or lumbar level, because the caudal regions of the vertebral column develop later.^6,7^ Abnormal Homeobox (*HOX)* gene expression is thought to be a causal factor.^1^ *HOX* genes are a group of highly conserved genes that have an important function in various developmental processes, including anteroposterior patterning and determination of vertebral identity.^15-17^ Many experiments on mice and chicks have shown that changed expression of specific *HOX* genes leads to homeotic transformations of vertebrae and abnormalities in different organ systems.^16,18-20^ Altered *HOX* gene expression has also been associated with the development of malignancies.^21,22^

The aims of the current study were to examine the prevalence of abnormal vertebral patterns in deceased fetuses and neonates and to determine whether an abnormal vertebral pattern is associated with (specific) structural and/or chromosomal anomalies in this population. This study also aimed to gain insight into the possible impact of DNA Copy Number Variations (CNVs) in coding regions and/or disturbing enhancers on the development of the pattern of the vertebral column.

## Methods

### Study population

The cohort consisted of fetuses and infants, younger than one year old, deceased between 2009 and 2015 in the Erasmus University Medical Center Sophia Children’s Hospital, of whom a babygram, autopsy report and/or SNP Array was available. Spontaneous intrauterine fetal demise and medically indicated terminations of pregnancies were included. Medical information about mothers, fetuses and neonates were retrieved from electronic medical records.

Babygrams were made routinely if autopsy was performed. Autopsies were performed according to the national guidelines.^23^ Congenital anomalies were categorized according to the European Registry of Congenital Anomalies and Twins (Eurocat) classification system.^24^ If autopsy had not been performed, the presence of malformations was based on the report of the prenatal advanced ultrasound scan, pre- or postnatal radiographic investigations or post-mortem external inspection. If none of these investigations had been requested, or the results were inconclusive (e.g. due to maceration), the presence of congenital anomalies was categorized as non-available.

Radiographs were made both ventrally and laterally (5.6-12.6mAs, 40-50 kV, Philips Optimus ZBM3/NZR91, Philips Medical Systems, Eindhoven, the Netherlands). Radiographs performed at other hospitals were requested. All radiographs were assessed by one reviewer, who was blinded for the autopsy results and the results of genetic investigations. Rudimentary ribs and the vertebral pattern were defined following ten Broek *et al.*^*6*^: if a rib on the most cranial or most caudal thoracic vertebra had a length of less than half of the rib of the adjacent thoracic vertebra, it was considered rudimentary. Rudimentary cervical ribs were scored if the length of the transverse processes of the seventh cervical vertebra was more than the transverse process of the first thoracic vertebra, but less than half of the first thoracic rib. If the rib was longer than half of the first thoracic rib, it was considered a (complete) cervical rib. Deviations from the normal vertebral pattern were classified as more severe when cranially located vertebral regions and multiple vertebral regions were involved (figure 1).

**Figure 1.**
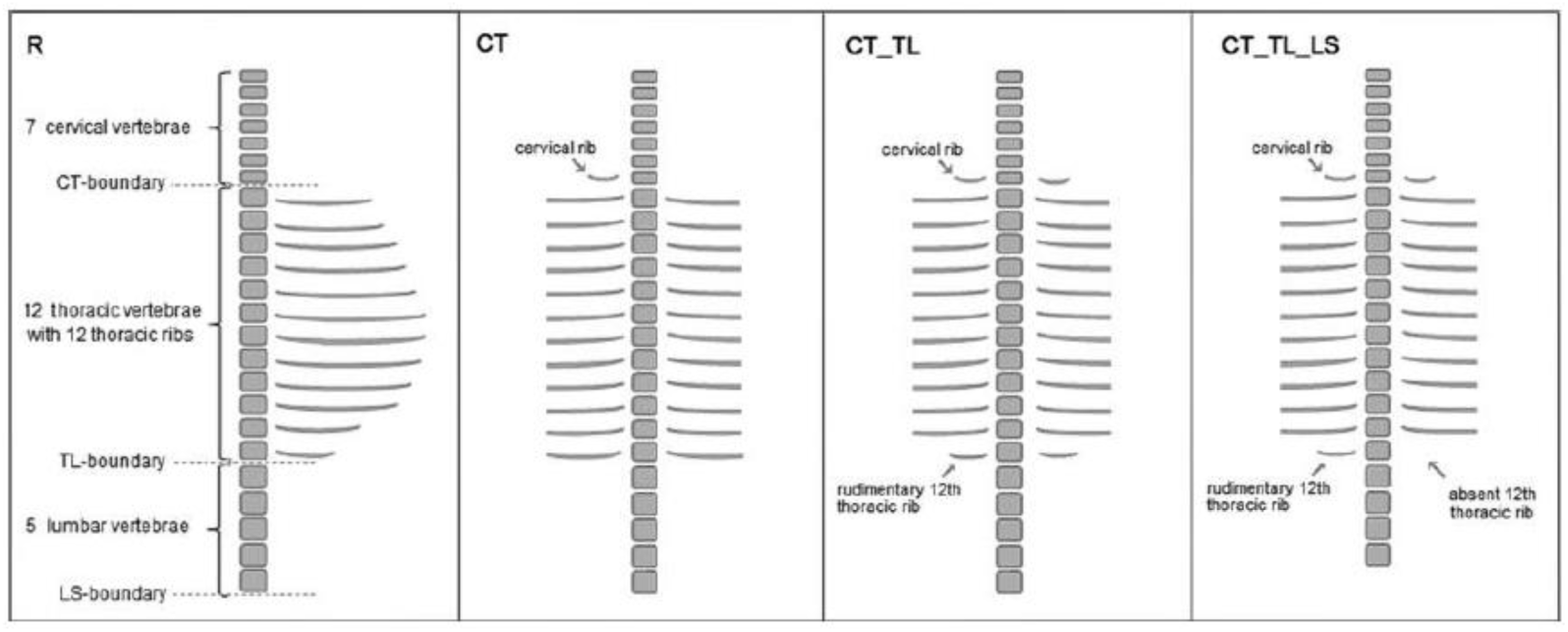
Overview of different vertebral columns and number on severity scale. From left to right: R= Regular pattern, severity scale value 0; CT= shift at the cervicothoracic boundary, severity scale value 4; CT_TL= shift at the cervicothoracic and thoracolumbar boundary, severity scale value 6; CT_TL_LS= shift at cervicothoracic, thoracolumbar and lumbosacral boundary, severity scale value 7.

A subset of thirty randomly chosen radiographs was assessed twice by the same reviewer and by a second reviewer, to determine the intraobserver and interobserver variability.

The study was performed according to the Dutch law on clinical trials in the Netherlands (WMO). Under this law the study does not require ethical approval or informed consent. This was confirmed by the institutional ethics committee (Medical Research Ethics Committee Erasmus MC, MEC-2014-098).

### Analysis of copy number variation

DNA was isolated from material that was collected in patients opting for invasive prenatal or postnatal diagnostic tests. We determined the CNV profiles in all coding and non-coding regions of patients (n=265) using methods and analysis settings previously described.^25^ CNV profiles were inspected visually in Biodiscovery Nexus CN8.0. (Biodiscovery Inc., Hawthorne, CA, USA). Rare CNVs were classified as described in table 1 and inspected for overlap with candidate genes, such as HOX genes and genes previously reported to be associated with vertebral anomalies in VACTER-L patients. ^26,27^ A more detailed description of this analysis can be found in the supplementary methods.

**Table 1.** Rare CNVs in prenatal cohort.

| Copy Number Loss |  | Number of cases |
| --- | --- | --- |
| Deleterious | ≥2Mb# | 10 |
| Likely deleterious | ≥1Mb | 6 |
| VOUS | ≤ 1Mb inheritance unspecified | 20 |
| Likely Benign | present in DDD controls | 7 |
|  |  | 43 |
| Copy Number Gain |  |  |
| Deleterious | ≥3Mb or overlapping dominant disease genes | 4 |
| Likely deleterious | ≥2Mb | 3 |
| VOUS | ≤ 1Mb inheritance unspecified | 17 |
| Likely Benign | present in DDD controls | 5 |
|  |  | 29 |

### Statistical analysis

Statistical analysis was performed using SPSS Statistics (IBM Corp. Released 2013. IBM SPSS Statistics for Windows, Version 21.0. Armonk, NY: IBM Corp.). Determination of differences between groups were calculated using independent sample t-tests for continuous data and chi-square test and Fisher’s exact test for categorical data. Descriptive statistics were used to evaluate outcome parameters. A p-value <0.05 was considered statistically significant. Kappa’s test was performed in order to determine the interobserver and intraobserver reliability. Kappa values between 0.61 and 0.80 were considered to be substantial and kappa values between 0.81 and 1.00 were considered almost perfect.^28^

## Results

The study population consisted of 374 fetuses and 71 neonates. In 71 fetuses and neonates (16.0%), the vertebral pattern could not be reliably assessed by radiography. This was due to inadequate positioning of the fetus or neonate, overprojection of the maxilla, clavicles or umbilical cord clamp, or low quality of the babygram. Excluding these 71 patients, 316 fetuses and 58 neonates were available for analysis of the pattern of the vertebral column. No statistically significant differences were found between the included and excluded patients in gestational age, presence or type of congenital anomaly and pregnancy outcome (table S1). Of the 374 included fetuses and neonates, 188 (50.3%) pregnancies were terminated because of medical reasons, mostly (suspected) fetal structural, chromosomal or other genetic anomalies (N=180/188, 95.7%). Other medical indications for pregnancy terminations were severe fetal growth restriction (N=1, 0.5%), preterm premature rupture of membranes (N=3, 1.6%) or poor maternal condition in pregnancies with early onset severe pre-eclampsia or HELLP syndrome (N=4, 2.1%). The second largest group consisted of miscarriages or intrauterine fetal demises (128/374, 34.2%). The majority of the 58 neonatal deaths was related to structural anomalies (43/58, 74.1%). Most other causes of neonatal death were associated with prematurity (table S2).

An overview of maternal, fetal and neonatal characteristics is provided in table 2. The inclusion of multiple pregnancies and more than one pregnancy per mother resulted in a total number of 366 included mothers. Autopsy was performed in the majority of patients (N=305, 81.6%). In 35 of the 69 patients in whom no autopsy was performed, an advanced ultrasound examination was carried out (9.4%). The presence of structural abnormalities could not be ascertained in 40 patients. This included 31 still births, 6 terminations of pregnancy and 3 live births. In 33 of these patients, neither autopsy nor advanced ultrasound examination had been performed. In 7 patients, the results of autopsy or advanced ultrasound examination were inconclusive.

**Table 2.**
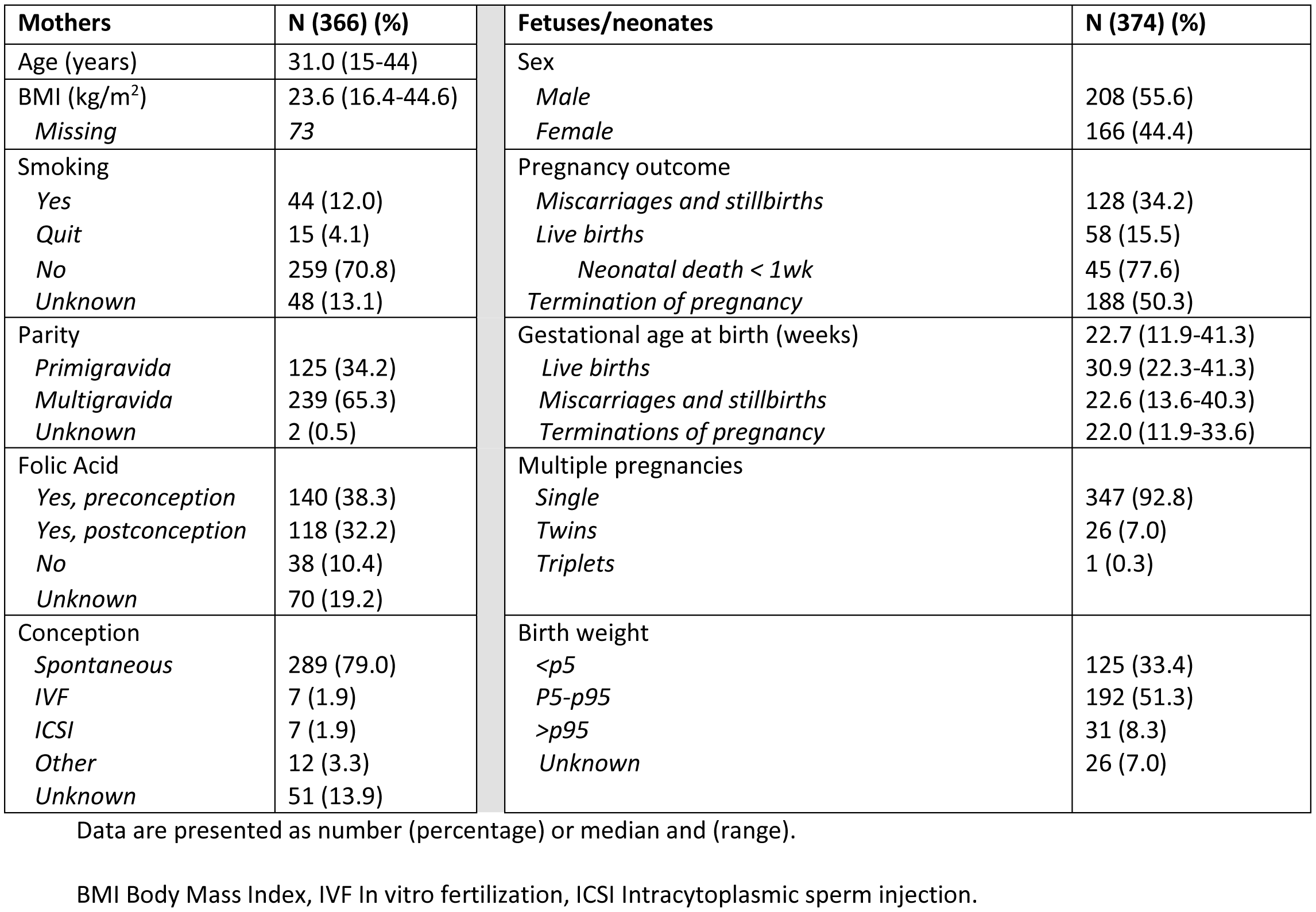
Characteristics of mothers, fetuses and neonates with a known vertebral pattern of the fetuses and neonates.

Structural anomalies were present in a large proportion of the group (256/334, 76.6%) and the prevalence was highest in the subgroup of pregnancy terminations (173/182, 95.1%), compared to 43/55 (78.2%) in live births and 40/97 (41.2%) in stillbirths. In more than half of the patients with a structural anomaly, more than one organ system was affected (146/256, 57.0%). The most frequently affected organ systems were the cardiovascular (N = 90), nervous (N = 85), craniofacial (N = 82), limbs (N = 71) and urogenital (N = 66) system (figure 2).

**Figure 2.**
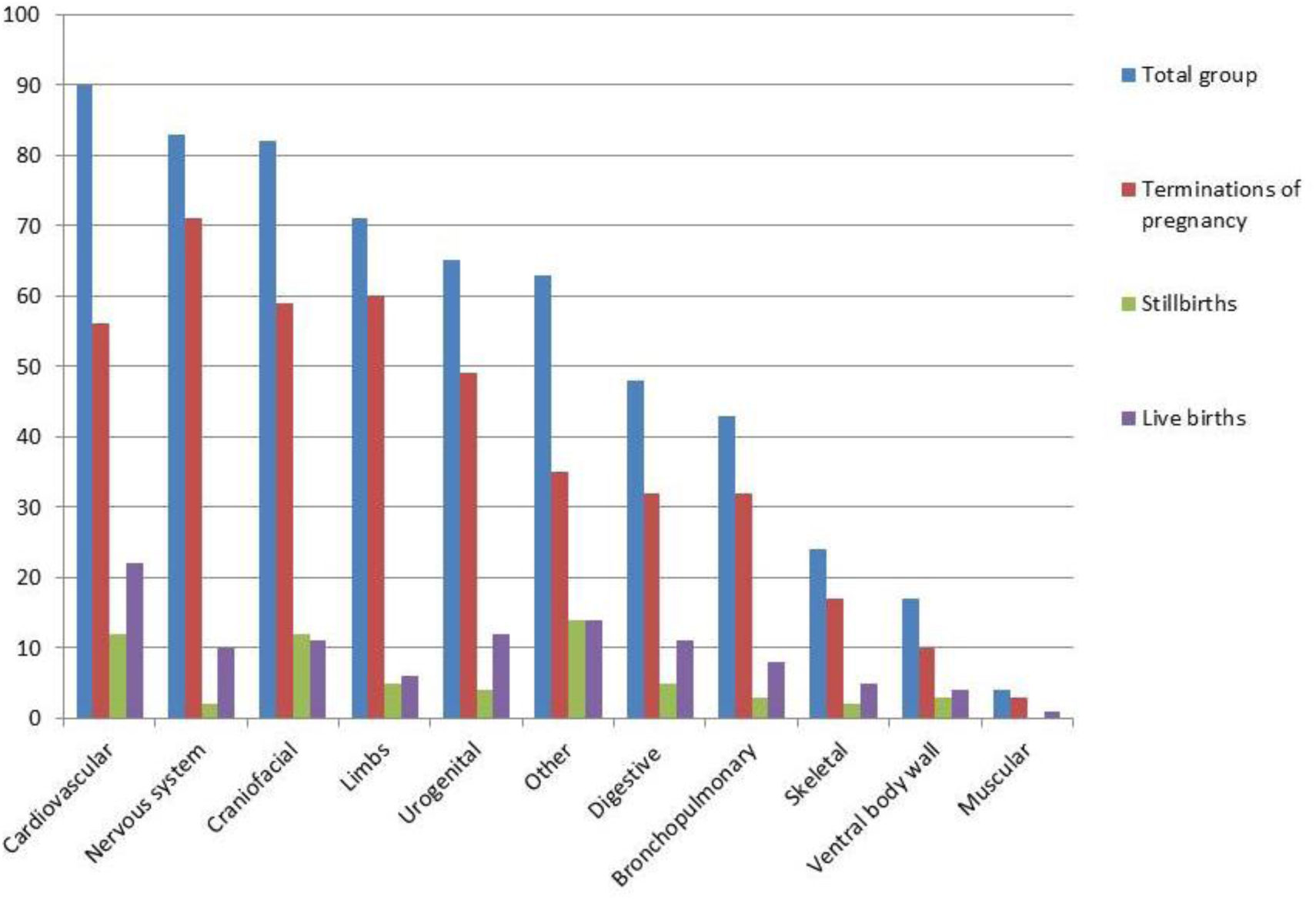
The distribution of structural abnormalities in the total study population and subgroups of live births, stillbirths and terminations of pregnancy. CT= shift at the cervicothoracic boundary, CT_TL= shift at the cervicothoracic and thoracolumbar boundary, CT_LS= shift at cervicothoracic and lumbosacral boundary, TL_LS= shift at thoracolumbar and lumbosacral boundary, CT_TL_LS= shift at cervicothoracic, thoracolumbar and lumbosacral boundary.

A regular vertebral pattern was identified in approximately one quarter of patients (100/374, 26.7%, figure 3). Of the 274 patients (73.3%) with an abnormal vertebral pattern, the cervicothoracic region was most often affected (N = 195/274, 71.2%), either exclusively, or in combination with thoracolumbar and/or lumbosacral shifts (CT (N = 80, 21.4%), CT_LS (N = 15, 4.0%), CT_TL (N = 84, 22.5%), CT_TL_LS (N = 16, 4.3%)). Cervical ribs were seen in approximately half of the patients (188/374, 50.3%). In the majority, cervical ribs were bilateral (N=128, 68.1%); of the unilateral cervical ribs, most were left sided (N=37/60, 61.7%). Figure 4 shows a babygram of a fetus with rudimentary twelfth thoracic ribs and rudimentary cervical ribs, and a babygram of a neonate with a regular vertebral pattern.

**Figure 3.**
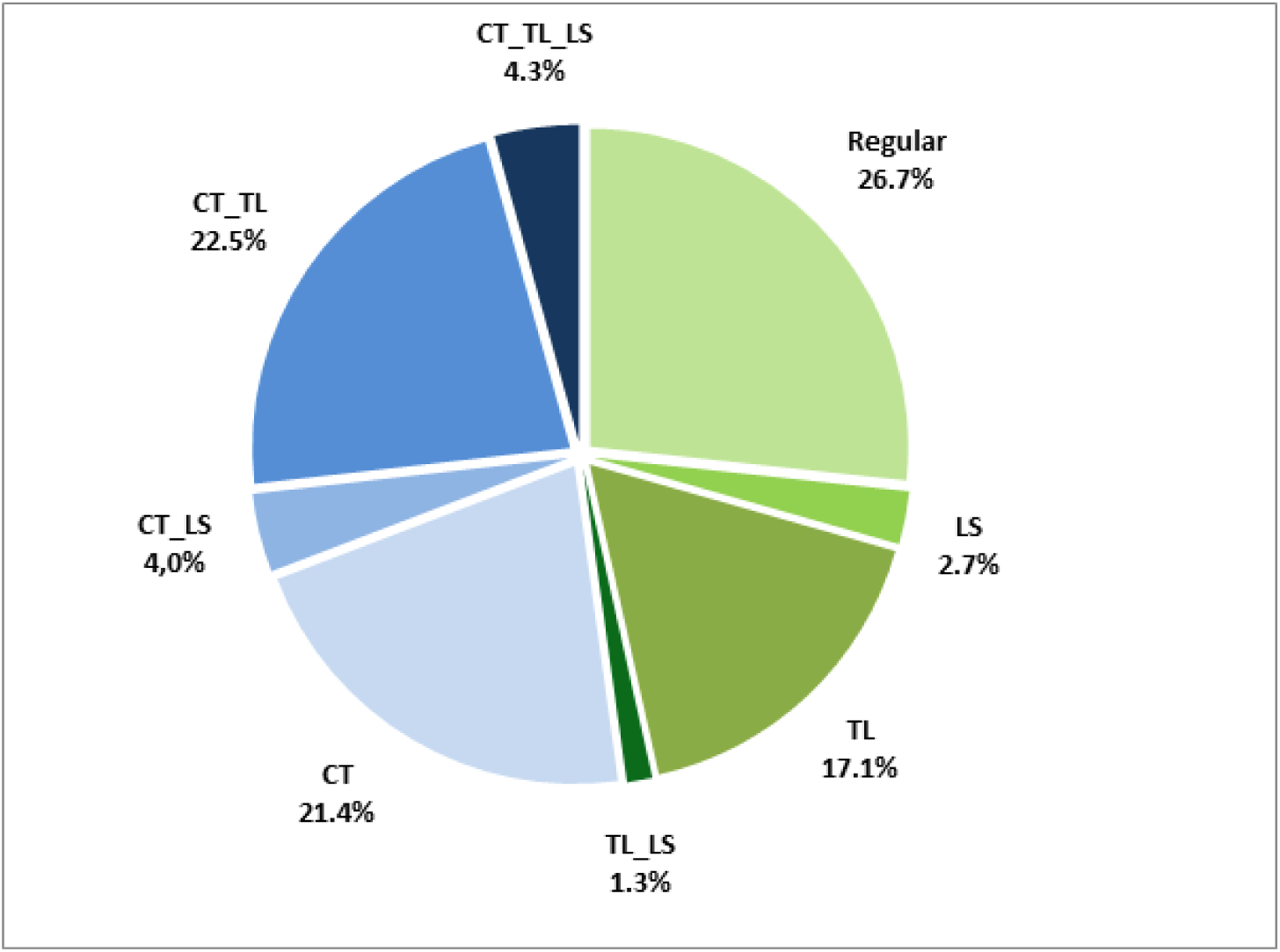
The distribution of the different patterns of the vertebral column in the total study population (N=374) CT= shift at the cervicothoracic boundary, CT_TL= shift at the cervicothoracic and thoracolumbar boundary, CT_LS= shift at cervicothoracic and lumbosacral boundary, TL_LS= shift at thoracolumbar and lumbosacral boundary, CT_TL_LS= shift at cervicothoracic, thoracolumbar and lumbosacral boundary.

**Figure 4.**
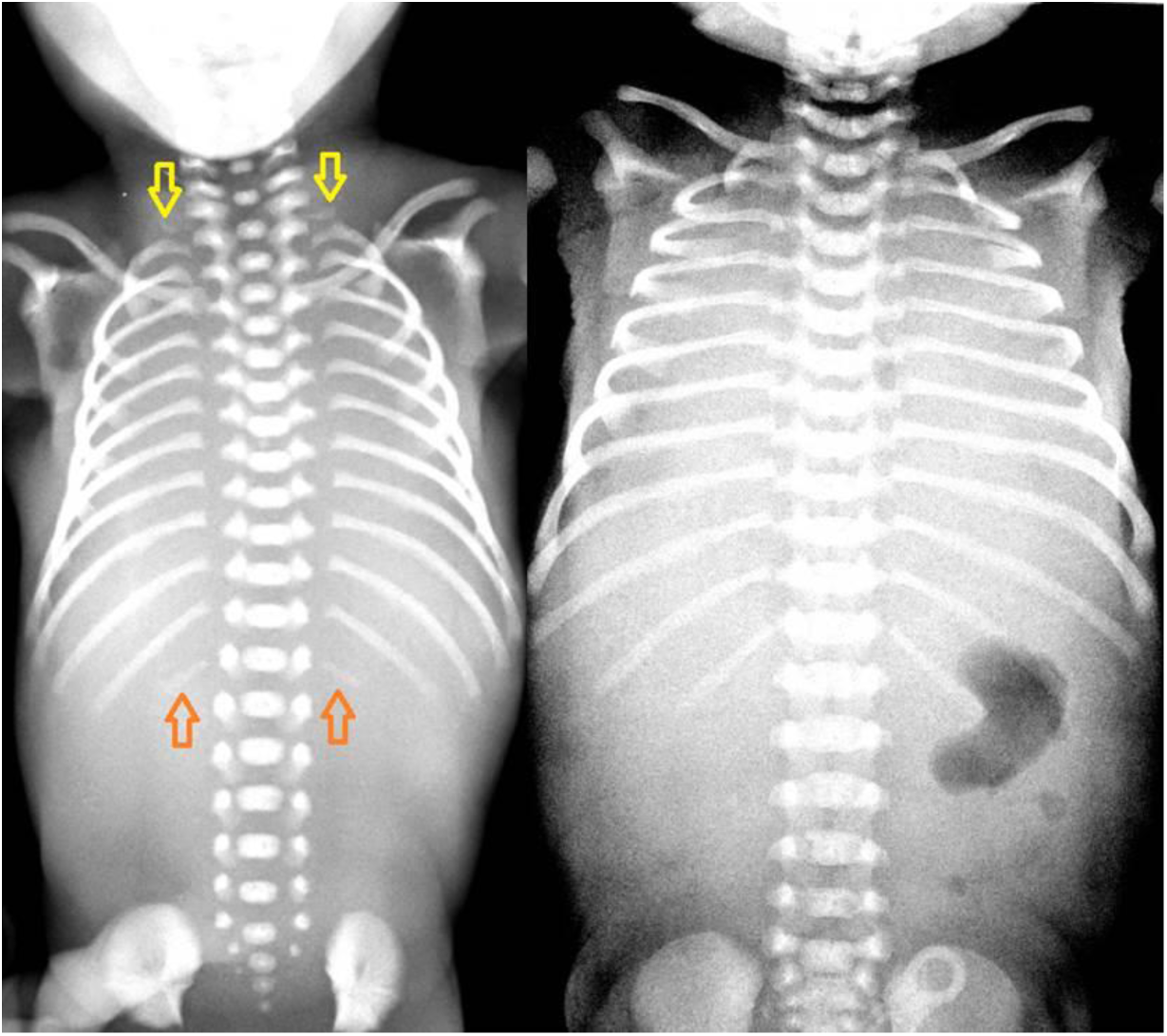
Babygram of a fetus at 20.1 weeks’ gestation (left) and a deceased term neonate (right). The babygram on the left shows rudimentary 12th thoracic ribs (orange arrows) and the presence of rudimentary cervical ribs (yellow arrows); the babygram on the right shows 12 thoracic rib pairs and the absence of cervical ribs.

The distribution of the vertebral pattern categorized according to structural abnormality is shown in figure 5. The prevalence of a regular vertebral pattern ranged between 18.6 and 33.3% and was highest in the group without structural abnormalities (26/78) and lowest in the group with bronchopulmonary abnormalities (8/43, 18.6%). Changes involving the cervicothoracic region occurred most frequently in patients with skeletal abnormalities (17/24, 70.8%), abnormalities involving the digestive system (28/46, 60.9%) and limb defects (43/71, 60.6%). The most disturbed vertebral pattern (CT_TL_LS) was frequently seen in patients with ventral body wall defects (3/17, 17.6%), skeletal (3/24, 12.5%) and craniofacial (9/82, 11.0%) abnormalities.

**Figure 5.**
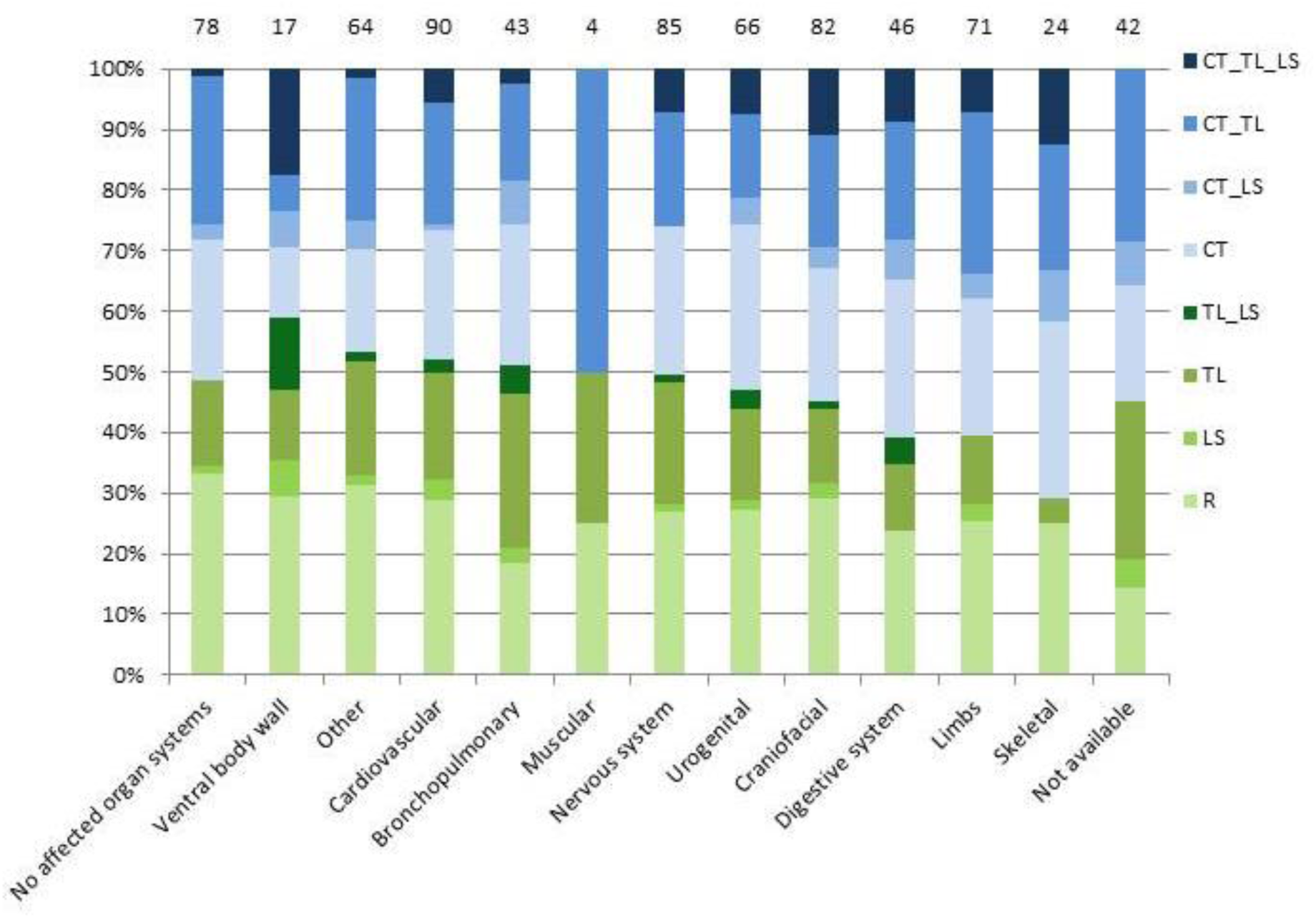
The pattern of the vertebral column in fetuses and neonates categorized according to the presence and type of structural anomaly. The total number of cases in each group is depicted above the bars. R= Regular pattern, CT= shift at the cervicothoracic boundary, CT_TL= shift at the cervicothoracic and thoracolumbar boundary, CT_LS= shift at cervicothoracic and lumbosacral boundary, TL_LS= shift at thoracolumbar and lumbosacral boundary, CT_TL_LS= shift at cervicothoracic, thoracolumbar and lumbosacral boundary.

After subdivision of the study population in stillbirths, live births and terminations of pregnancies, it became clear that the proportion of fetuses and neonates with a regular vertebral pattern was significantly higher in live births (25/58, 43.1%), compared to stillbirths (29/128, 22.7%) and terminations of pregnancies (46/188, 24.5%, p = 0.009). Cervical ribs were significantly more common in stillbirths (69/128 (53.9%) and terminations of pregnancies (101/188 (53.7%), compared to live births (18/58, 31.0%, p = 0.006). The distribution of the vertebral pattern in these subgroups is shown in figure 6. The prevalence of cervical ribs did not differ significantly between fetuses and neonates with and without structural anomalies in the total group (126/256 versus 40/78, X^2^ (1) = 0.10, p = 0.80), nor in the subgroups of live births (13/43 versus 4/12, p = 1.0), stillbirths (21/40 versus 32/57, X^2^ (1) = 0.13, p = 0.72), or terminations of pregnancies (92/173, versus 4/9, p = 0.74). The 4 live births with cervical ribs, but without structural abnormalities died because of a subgaleal hemorrhage, sepsis, asphyxia and uterine rupture, respectively. Autopsy was performed in all of these neonates. The asphyctic neonate had mild dysmorphic features, but a normal karyotype. The pregnancy in which a uterine rupture occurred was complicated by polyhydramnios and macrosomia.

**Figure 6.**
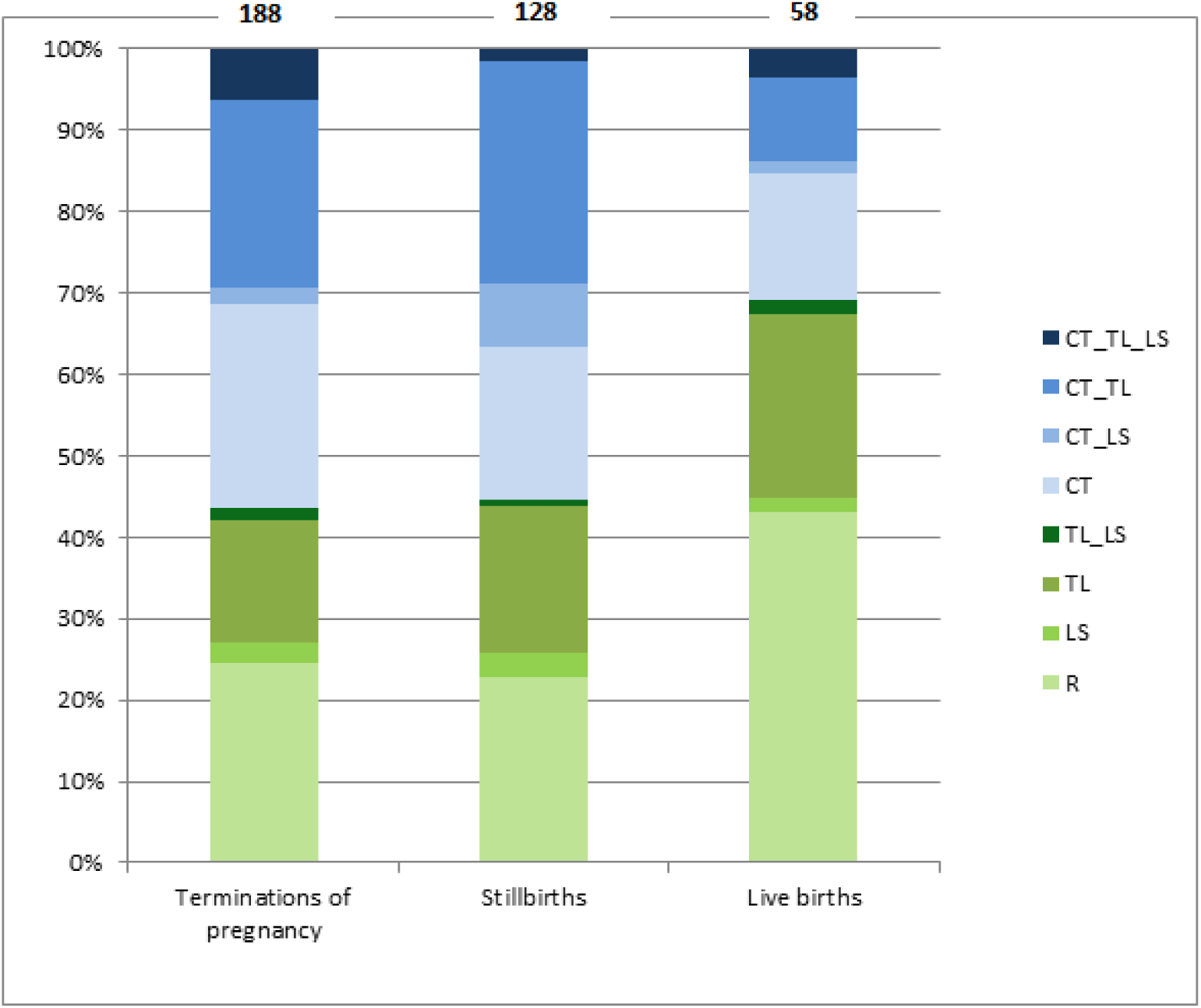
The distribution of the vertebral pattern in terminations of pregnancies, still births and live births. The total number of cases is depicted above the bars. R= Regular pattern, CT= shift at the cervicothoracic boundary, CT_TL= shift at the cervicothoracic and thoracolumbar boundary, CT_LS= shift at cervicothoracic and lumbosacral boundary, TL_LS= shift at thoracolumbar and lumbosacral boundary, CT_TL_LS= shift at cervicothoracic, thoracolumbar and lumbosacral boundary.

Maternal diabetes was ruled out. The macrosomic neonate had mild dysmorphic features and was suspected of having an overgrowth syndrome, but additional DNA-testing did not reveal a genetic mutation. Chromosomal or genetic analyses were not performed in the remaining 2 neonates.

No statistically significant difference was found in the prevalence of cervical ribs between males and females (109/208 versus 79/166, X^2^ (1) = 0.86, p = 0.36). When the group was categorized according to the number of affected organ systems, the proportion of fetuses and neonates with a disturbed vertebral pattern involving all vertebral boundaries (CT_TL_LS) was highest in the subgroup with four or more affected organ systems (fig S1).

Gestational age at birth (M = 23.5 weeks, SD = 6.2, versus M = 23.7 weeks, SD=6.3, p = 0.73), maternal smoking (p = 0.39), assisted conception (p = 0.47), use of folic acid (p = 0.77) and maternal BMI (M = 24.8, SD = 5.3, versus M = 24.7, SD = 4.7, p = 0.91) were not significantly different between fetuses and neonates with and without cervical ribs.

The intra-observer reliability was almost perfect for determination of the number of thoracic ribs (Kappa = 0.84) and substantial for assessment of the presence of cervical ribs (Kappa = 0.78). The interrater reliability was substantial for both determination of the number of thoracic ribs (Kappa = 0.77) and cervical ribs (Kappa = 0.74).

Karyotype was available in 221 (59.1%) patients. Aneuploidies were detected in 15 (6.8%) patients; 10 (66.7%) of these had cervical ribs. Microarray tests were performed in 265 (70.9%) of patients. Neither karyotype nor array was available in 54 patients (14.4%). In 31 patients (31/265, 11.7%) microarray showed an abnormality. Cervical ribs were present in 20/31 (64.5%) patients with an abnormal microarray result. The presence of cervical ribs in patients with a pathogenic or (likely) deleterious CNV was 50.0% (6/12) and 57.1% (4/7), respectively. These data are summarized by subgroup in table S3.

### CN profiling

The prevalence of large (likely) deleterious CNV (table 2) and aneuploidies was high. However, there were no recurrent or overlapping rare CNVs. Some of the rare CNVs have overlap with dominant disease genes (see table S4). There was no overlap with candidate genes (e.g. the *HOX* gene cluster) Pathway enrichment analysis of genes affected by a rare CNV did not provide evidence of enrichment of relevant pathways.

## Discussion

An abnormal pattern of the vertebral column, including the presence of cervical ribs, was frequently found in deceased fetuses and neonates. The prevalence was noticeably higher in this population, compared to living children or adults who are not known to have structural, chromosomal or genetic abnormalities, as reported in the literature.^9^ These findings are in line with the study of ten Broek et al., who included a similar study population.^6^ The differences in vertebral pattern between subgroups categorized into affected organ system were less obvious compared to the study of ten Broek et al.^6^, which could be due to the smaller number of included patients in our study. The high frequency of the most severely disturbed vertebral pattern in the subgroups with ventral body wall defects, craniofacial and skeletal malformations was confirmed. The co-occurrence of an abnormal vertebral pattern and these abnormalities could be explained by the close spatial relationship and the intense interaction of signaling pathways between the embryonic precursors of these organ systems and the vertebral column in early embryogenesis.^29-31^ For instance, signaling from the somites influences the migration of neural crest cells, from which the craniofacial skeleton is derived, whereas somites themselves give rise to the vertebrae and ribs, ventral body wall skeleton, abdominal muscles, skeletal muscles and cartilage. Because of the intense interactions between developmental processes during somitogenesis, this period is considered an extremely vulnerable period.^32,33^ Consequently, disruptions in vertebral patterning would often be accompanied by disruptions of other developmental processes. As expected, the most severely disturbed vertebral pattern was most frequent in patients with more than four affected organ systems. This might be due to a prolonged disturbance of developmental processes.

Unexpectedly, the prevalence of cervical ribs was not significantly different between fetuses and neonates with and without structural abnormalities. This is in contrast to ten Broek et al.^6^, but has also been observed by Furtado et al.^14^, who suggested that incomplete penetrance of genetic events leading to the occurrence of both abnormal vertebral patterning and fetal death, could result in a milder phenotype without structural abnormalities. Thus the presence of cervical ribs might not only be associated with the presence of structural abnormalities, but also with the occurrence of intrauterine fetal demise itself, even in the absence of structural abnormalities detected by autopsy. The significantly higher prevalence of cervical ribs in the subpopulation of stillbirths compared to live births is in line with this theory. The relatively high prevalence of cervical ribs within the subpopulation of deceased neonates without proven structural abnormalities (4/11, 36.4%) remains unexplained, however. The disappearance of cervical ribs later in fetal life or childhood has also been considered, but seems less plausible, as cervical ribs are frequently encountered in specific (adult) patient groups.^11,12,34^ In addition, no differences between the presence of cervical ribs in fetuses and infants was found by Galis et al.^7^ and the prevalence of cervical ribs was not substantially lower in adults compared to children in the general population.^9^

Cervical ribs were detected in the majority of patients with an aneuploidy or abnormal microarray result. The reported prevalence of cervical ribs in aneuploidies ranges between 12.5 and 100%.^14,35^ Studies reporting on cervical ribs in populations with microarray abnormalities are lacking. The number of patients with similar chromosomal or genetic abnormalities in this cohort was insufficient to enable the detection of statistically significant associations between cervical ribs and specific chromosomal or genetic abnormalities.

Overlapping abnormalities involving *HOX*-genes or other candidate genes were not identified within this cohort. This does not completely rule out a common genetic basis for the abnormalities of the vertebral pattern, because the presence of structural DNA variations, such as point mutations, are not detected by SNP array and CNV analyses. However, both the absence of rare recurrent CNVs involving candidate genes, and the fact that several patients included in this study population proved to have different chromosomal or genetic abnormalities are an indication of the genetic heterogeneity that appears to underlie the development of an abnormal vertebral pattern. In addition, it is unlikely that half of the stillbirths (those with cervical ribs) had a common underlying genetic basis.

The occurrence of similar segmentation anomalies in patients with various structural, chromosomal and genetic abnormalities and prenatal exposure to different teratogens, seems to be a reflection of the different underlying etiologies of vertebral homeotic transformations.^36,37^ This heterogeneity may be explained by the intense interactivity during the head-to-tail patterning of the cervical vertebrae, such that many disturbances can disrupt this patterning. The specific timing and duration of the disruption has greater influence than the cause of the disruption itself.^38-44^ Abnormalities in vertebral patterning, such as cervical ribs, could thus be regarded as a sign of abnormal embryonic development, irrespective of the causative event. The remarkably high frequency of cervical ribs and other deviations from the regular pattern of the vertebral column in this study population of deceased fetuses and neonates, underwrites the hypothesized selection against variations in the conserved process of vertebral patterning.

These findings indicate that assessment of the vertebral pattern could be of added value in determination of fetal and neonatal outcome. Prenatal assessment of the vertebral pattern, including the detection of rudimentary cervical ribs, using (3-dimensional) ultrasound seems feasible.^45-50^

Strengths of this study are the large study population and the good intraobserver and interobserver reliability for the assessment of the vertebral pattern and cervical ribs on radiographs. Limitations are the small sizes of the subgroups with specific structural, chromosomal or genetic abnormalities and the fact that autopsy had not been performed in all patients. Although the presence of a healthy control group was lacking, literature regarding the prevalence of cervical ribs in healthy populations was available for comparison.

## Conclusion

The presence of abnormalities in the pattern of the vertebral column, particularly in the cervical region, could be regarded as a sign of disruption at critical, interactive and conserved stages of early embryonic development. The absence of rare recurrent CNVs, and the presence of similar vertebral patterning abnormalities in patients with different chromosomal or genetic abnormalities are an indication of the genetic heterogeneity that appears to underlie the development of an abnormal vertebral pattern. Assessment of the vertebral pattern could provide valuable information regarding fetal and neonatal outcome.

Further studies investigating the feasibility and value of prenatal (3-dimensional) ultrasound assessment of the number of vertebrae and ribs are warranted. Whole exome sequencing on subjects with (isolated) vertebral patterning abnormalities might provide insight into the presumably heterogeneous genetic causes of these patterning defects.

## Acknowledgements

The authors thank Ms. S.C. Husen^a^, for the assessment of the subset of radiographs for determination of the interobserver reliability. No funding source or compensation.

## Data availability

The datasets generated during and/or analyzed during the current study are available from the corresponding author on reasonable request.

## References

1 Galis, F. Why do almost all mammals have seven cervical vertebrae? Developmental constraints, Hox genes, and cancer. J Exp Zool 285, 19-26 (1999).

2 Brewin, J., Hill, M. & Ellis, H. The prevalence of cervical ribs in a London population. Clin Anat 22, 331-336 (2009).

3 Etter, M. Osseous abnormalities of the thoracic cage seen in forty thousand consecutive chest photoroentgenograms. Am J Roentgenol 51, 359-363 (1944).

4 Bots, J. et al. Analysis of cervical ribs in a series of human fetuses. J Anat 219, 403-409 (2011).

5 Narita, Y. & Kuratani, S. Evolution of the vertebral formulae in mammals: a perspective on developmental constraints. J exp zool B Mol Dev Evol 304, 91-106 (2005).

6 Ten Broek, C. M. et al. Evo-Devo of the Human Vertebral Column: On Homeotic Transformations, Pathologies and Prenatal Selection. Evol Biol 39, 456-471 (2012).

7 Galis, F. et al. Extreme selection in humans against homeotic transformations of cervical vertebrae. Evolution 60, 2643-2654 (2006).

8 Varela-Lasheras, I. et al. Breaking evolutionary and pleiotropic constraints in mammals: On sloths, manatees and homeotic mutations. Evodevo 2, 11 (2011).

9 Schut P. C. et al. Adverse Fetal and Neonatal Outcome and an Abnormal Vertebral Pattern: A Systematic Review. Obstetrical & Gynecological Survey 71, 741-750 (2016).

10 Loder, R. T., Huffman, G., Toney, E., Wurtz, L. D. & Fallon, R. Abnormal rib number in childhood malignancy: Implications for the scoliosis surgeon. Spine 32, 904-910 (2007).

11 Zierhut, H., Murati, M., Holm, T., Hoggard, E. & Spector, L. G. Association of rib anomalies and childhood cancers. Br J Cancer 105, 1392-1395 (2011).

12 Merks, J. H. M. et al. Prevalence of RIB anomalies in normal Caucasian children and childhood cancer patients. Eur J Med Genet 48, 113-129 (2005).

13 Schumacher, R., Mai, A. & Gutjahr, P. Association of rib anomalies and malignancy in childhood. Eur J Pediatr 151, 432-434 (1992).

14 Furtado, L. V., Thaker, H. M., Erickson, L. K., Shirts, B. H. & Opitz, J. M. Cervical ribs are more prevalent in stillborn fetuses than in live-born infants and are strongly associated with fetal aneuploidy. Pediatr Dev Pathol 14, 431-437 (2011).

15 Lappin, T. R., Grier, D. G., Thompson, A. & Halliday, H. L. HOX genes: seductive science, mysterious mechanisms. Ulster Med J 75, 23-31 (2006).

16 Mallo, M., Wellik, D. M. & Deschamps, J. Hox genes and regional patterning of the vertebrate body plan. Dev Biol 344, 7-15 (2010).

17 Quinonez, S. C. & Innis, J. W. Human HOX gene disorders. Mol Genet Metab 111, 4-15 (2014).

18 Horan, G. S., Wu, K., Wolgemuth, D. J. & Behringer, R. R. Homeotic transformation of cervical vertebrae in Hoxa-4 mutant mice. Proc Natl Acad Sci U S A 91, 12644-12648 (1994).

19 Wellik, D. M., Hawkes, P. J. & Capecchi, M. R. Hox11 paralogous genes are essential for metanephric kidney induction. Genes Dev 16, 1423-1432 (2002).

20 Manley, N. R. & Capecchi, M. R. Hox group 3 paralogs regulate the development and migration of the thymus, thyroid, and parathyroid glands. Dev Biol 195, 1-15 (1998).

21 Anbazhagan, R., Raman V.. Homeobox genes: molecular link between congenital anomalies and cancer. Eur J Cancer 33, 635-637 (1997).

22 van der Lugt, N. M. et al. Posterior transformation, neurological abnormalities, and severe hematopoietic defects in mice with a targeted deletion of the bmi-1 proto-oncogene. Genes Dev 8, 757-769 (1994).

23 https://www.nvk.nl/Nieuws/Dossiers/NODO. (September 12th 2018).

24 EUROCAT. Coding of EUROCAT Subgroups of Congenital Anomalies. Chapter 3.3, Guide 1.4 (2012).

25 Brosens, E. et al. Copy number variations in 375 patients with oesophageal atresia and/or tracheoesophageal fistula. Eur J Hum Genet 24, 1715-1723 (2016).

26 Chen, Y. et al. The genetic landscape and clinical implications of vertebral anomalies in VACTERL association. J Med Genet 53, 431-437, doi:jmedgenet-2015-103554 [pii]10.1136/jmedgenet-2015-103554 (2016).

27 Solomon, B. D. et al. Clinical geneticists’ views of VACTERL/VATER association. Am J Med Genet A 158A, 3087-3100 (2012).

28 Landis, J. R. & Koch, G. G. The measurement of observer agreement for categorical data. Biometrics 33, 159-174 (1977).

29 Mekonen, H. K., Hikspoors, J. P., Mommen, G., Kohler, S. E. & Lamers, W. H. Development of the ventral body wall in the human embryo. J Anat 227, 673-685 (2015).

30 Rajion, Z. A. et al. A three-dimensional computed tomographic analysis of the cervical spine in unoperated infants with cleft lip and palate. Cleft Palate Craniofac J 43, 513-518 (2006).

31 Sonnesen, L. Associations between the Cervical Vertebral Column and Craniofacial Morphology. Int J Dent 2010, 295728 (2010).

32 Lubinsky, M. Blastogenetic associations: General considerations. Am J Med Genet A 167A, 2589-2593 (2015).

33 Galis, F. & Metz, J. A. Testing the vulnerability of the phylotypic stage: on modularity and evolutionary conservation. J Exp Zool 291, 195-204, doi:10.1002/jez.1069 (2001).

34 Weber, A. E. & Criado, E. Relevance of bone anomalies in patients with thoracic outlet syndrome. Ann Vasc Surg 28, 924-932 (2014).

35 Schut, P. C. et al. Increased prevalence of abnormal vertebral patterning in fetuses and neonates with trisomy 21. J Matern Fetal Neonatal Med, 1-7 (2018).

36 Martinez-Frias, M. L. Segmentation anomalies of the vertebras and ribs: one expression of the primary developmental field. Am J Med Genet A 128A, 127-131, doi:10.1002/ajmg.a.30016 (2004).

37 Giampietro, P. F. et al. Clinical, genetic and environmental factors associated with congenital vertebral malformations. Mol Syndromol 4, 94-105 (2013).

38 JG, W. Methods for administering agents and detecting malformations in experimental animals. In: Wilson, J.G. and Warkany, J., Eds., Teratology: Principals and Techniques University of Chicago Press, Chicago, 262-277 (1965).

39 Lu, C. C., Matsumoto, N. & Iijima, S. Teratogenic effects of nickel chloride on embryonic mice and its transfer to embryonic mice. Teratology 19, 137-142, doi:10.1002/tera.1420190202 (1979).

40 Lu, F. Basic toxicology. Fundamentals, target organs and risk assessment.. Bristol, PA: Taylor and Francis. (1991).

41 Sadler, T. W. Effects of maternal diabetes on early embryogenesis: II. Hyperglycemia-induced exencephaly. Teratology 21, 349-356, doi:10.1002/tera.1420210311 (1980).

42 De Sesso JM, H. S. Principles underlying developmental toxicity. In: Fan AM, Chang LW, editors. Toxicology and risk assessment principles, methods, and applications., 37-56 (1996).

43 Opitz, J. M. The developmental field concept. Am J Med Genet 21, 1-11, doi:10.1002/ajmg.1320210102 (1985).

44 Lubinsky, M. Associations in clinical genetics with a comment on the paper by Evans et al. on tracheal agenesis. AM J MED GENET 21, 35-38 (1985).

45 Khodair, S. A., Hassanen, O.A. Abnormalities of fetal rib number and associated fetal anomalies using three dimensional ultrasonography. The Egyptian Journal of Radiology and Nuclear Medicine 45, 689-694 (2014).

46 Hershkovitz, R. Prenatal diagnosis of isolated abnormal number of ribs. Ultrasound Obstet Gynecol 32, 506-509 (2008).

47 Gindes, L., Benoit, B., Pretorius, D. H. & Achiron, R. Abnormal number of fetal ribs on 3-dimensional ultrasonography: Associated anomalies and outcomes in 75 fetuses. J Ultrasound Med 27, 1263-1271 (2008).

48 T. Esser, P. R., N. Sarioglu, K.D. Kalache.. Three-dimensional ultrasonographic demonstration of agenesis of the 12th rib in a fetus with trisomy 21. Ultrasound in Obstetrics & Gynecology 27, 712-715 (2006).

49 A. Dall’Asta, G. P., C.C. Lees. Crystal Vue technique for imaging fetal spine and ribs. Ultrasound in Obstetrics & Gynecology 47, 383-384 (2016).

50 Schut, P.C., Verdijk, R. M., Joosten, M. & Eggink, A. J. Prenatal diagnosis of cervical ribs by three-dimensional ultrasound in a foetus with a herniated Dandy-Walker cyst. BMJ Case Rep 11, doi:11/1/e225381 [pii]10.1136/bcr-2018-225381 (2018).

