## Supplementary material for "“Abnormal vertebral patterns in genetically heterogeneous deceased fetuses and neonates: evidence of selection against variations”": Table S1-S4, figure S1, supplementary methods

**Table S1. Characteristics of the included fetuses and neonates in whom the vertebral pattern could be assessed and the excluded fetuses and neonates in whom the vertebral pattern could not be assessed.**

| | **Vertebral pattern assessable (n=374)** | **Vertebral pattern not assessable (n=71)** | **p-value** |
|---|---|---|---|
| Gestational age at birth (weeks) | 22.7 (11.9-41.3) | 21.4 (12.0-40.4) | 0.06 |
| Presence congenital anomaly | 256 (68.4) | 49 (69.0) | 0.36 |
| No congenital anomaly | 78 (20.9) | 11 (15.5) | |
| *Unknown* | 40 (10.7) | 11 (15.5) | |
| Type congenital anomaly | | | 0.24 |
| *Craniofacial* | 10 (3.9) | 0 | |
| *Nervous system* | 25 (9.8) | 3 (5.7) | |
| *Bronchopulmonary* | 4 (1.6) | 1 (1.9) | |
| *Cardiovascular* | 27 (10.6) | 5 (9.4) | |
| *Ventral body wall* | 3 (1.2) | 1 (1.9) | |
| *Digestive system* | 3 (1.2) | 0 | |
| *Urogenital* | 12 (4.7) | 6 (11.3) | |
| *Limb defects* | 9 (3.5) | 1 (1.9) | |
| *Skeletal* | 4 (1.6) | 4 (7.5) | |
| *Other* | 13 (5.1) | 3 (5.7) | |
| *Multiple* | 146 (57.3) | 29 (54.7) | |
| Pregnancy outcome | | | 0.79 |
| *Miscarriages and stillbirths* | 128 (34.2) | 25 (35.2) | |
| *Live births* | 58 (15.5) | 13 (18.3) | |
| *Termination of pregnancy* | 188 (50.3) | 33 (46.5) | |

Data are presented as number (percentage) or median and (range).

**Table S2. Gestational age at birth, age at death, presence of congenital abnormalities and cause of death of the included live births.**

| GA at birth (weeks) | Age at death (days) | Cervical rib(s) | Congenital abnormalities | Cause of death |
|---|---|---|---|---|
| 22.3 | 0 | No. | None. | Immature delivery. |
| 22.7 | 0 | Yes. | Transposition of the great arteries, dysmorphic facial features. | Immature delivery. |
| 23.6 | 0 | No. | None. | Immature delivery. |
| 23.9 | 0 | No. | Unknown (no autopsy, no advanced anomaly scan). | Immature delivery. |
| 24.0 | 0 | No. | Right isomerism left lung. | Unsuccessful resuscitation. |
| 24.4 | 0 | No. | Colpocephaly. | Palliative care because of extremely preterm birth. |
| 24.0 | 7 | No. | Mild retrognathia. | NEC and sepsis. |
| 24.6 | 7 | Yes. | None. | Sepsis and septic thrombus and emboli in multiple organs, leading to multiorgan dysfunction. NEC. |
| 24.6 | 12 | No. | Dysmorphic facial features. | Sepsis leading to cardiorespiratory failure. |
| 24.7 | 5 | No. | Unknown (no autopsy or advanced anomaly scan). | Bowel perforation, IRDS, discontinuation of treatment due to bowel ischemia. |
| 24.9 | 2 | No. | None. | Discontinuation of treatment because of cardiorespiratory failure and refractory lactic acidosis, possibly due to NEC. |
| 24.9 | 30 | No. | Ascites. | Circulatory failure, likely due to NEC. |
| 25.0 | 0 | No. | Bicuspid aortic valve. | Pulmonary hypoplasia (anhydramnios from GA 17 weeks) |
| 25.0 | 25 | No. | No malformations. | Respiratory and hemodynamic insufficiency. Sepsis. Right atrial thrombus. |
| 25.1 | 15 | No. | Hypertrophic left cardiac ventricle. | Sepsis, circulatory insufficiency, hypovolemic shock eci. |
| 26.1 | 0 | Yes. | Unknown (no autopsy, no advanced anomaly scan). | Emergency caesarian section because of placental abruption. Unsuccessful resuscitation. |
| 26.4 | 33 | No. | None. | Cardiorespiratory insufficiency and sepsis (NEC). |
| 27.4 | 0 | Yes. | Dysmorphic facial features, lateral deviation foot, atrial septal defect. | PPROM, tachycardia and suspected intrauterine infection, for which emergency caesarian section. |

| | | | | |
|---|---|---|---|---|
| | | | | Cardiorespiratory failure, unsuccessful resuscitation at birth. |
| 27.4 | 1 | No. | Hydrops, periventricular leukomalacia, left isomerism right lung, dysmorphic facial features. | Refractory cardiorespiratory failure in hydropic neonates with anemia and thrombocytopenia. |
| 27.7 | 0 | No. | None. | Cardiorespiratory failure due to septic shock (Klebsiella pneumoniae and NEC). |
| 27.9 | 1 | No. | Omphalocele, dysmorphic facial features, macroglossia, enlarged kidneys, open duct Botalli, periventricular leukomalacia. Beckwith Wiedemann syndrome. | Circulatory failure due to refractory hypotension. |
| 27.9 | 31 | No. | Ambiguous genital, macroglossia, short limbs. | Cardiorespiratory failure due to NEC. |
| 28 | 5 | No. | None. | Discontinuation of treatment because of severe asphyxia, pulmonary hypertension and immature lungs. |
| 29.0 | 0 | Yes. | No autopsy. Advanced anomaly scan and babygram suggestive of lethal skeletal dysplasia. | Palliative care. |
| 29.3 | 0 | No. | Omphalocele, hypertelorism, left-sided diaphragmatic hernia, hypoplastic lungs, ascites, hydronephrosis, dilated bladder. | Palliative care because of multiple congenital anomalies |
| 29.7 | 2 | No. | Hydrops, neuronal heterotopia. | Discontinuation of treatment because of poor prognosis due to extensive cystic periventricular leukomalacia. |
| 30.0 | 0 | Yes. | Pulmonary hypoplasia, polymicrogyria, glioneuronal heterotopia, skin lesions, ascites, hepatosplenomegaly, cholestasis, enlarged small bowel loops. | Respiratory failure. |

| | | | | |
|---|---|---|---|---|
| 30.7 | 0 | No. | Lissencephaly, hypoplastic corpus callosum, bicuspid aortic valve, pleural effusion. | Unsuccessful resuscitation after emergency caesarean section. |
| 30.7 | 0 | No. | Hydrops. Severe lung hypoplasia. | Respiratory insufficiency. |
| 31.0 | 0 | No. | Lissencephaly, neuronal heterotopia, ventriculomegaly, coarctation of the aorta, persistent left superior vena cava, ureteropelvic junction, dysmorphic facial features. | Discontinuation of treatment because of multiple abnormalities. |
| 31.0 | 1 | No. | Hydrops, hypertrophic dilated cardiomyopathy | Cardiac failure. |
| 32.0 | 6 | No. | Hydrops, transient myeloproliferative disorder, trisomy 21. | Discontinuation of treatment because of pulmonary hypertension not responsive to therapy. |
| 32.1 | 0 | Yes. | Pleural effusion, dysplastic aortic and pulmonary valves, ductus venosus agenesis, dysmorphic facial features, Noonan syndrome. | Cardiorespiratory failure. |
| 32.1 | 0 | No. | Suspicion of urethral valves. | Placental abruption, emergency caesarian section. Unsuccessful resuscitation. |
| 32.3 | 0 | No. | No autopsy. Advanced anomaly scan and prenatal MRI: right diaphragmatic hernia, Dandy Walker malformation, ventriculomegaly, persistent left superior vena cava. | Palliative care because of multiple abnormalities. |
| 32.3 | 1 | No. | Hydrops. | IRDS. |
| 32.7 | 0 | Yes. | Macrosomia. | Uterine rupture. Unsuccessful resuscitation. |
| 33.6 | 22 | No. | Pontocerebellar dysplasia. | Respiratory failure due to central apnea. |
| 33.7 | 9 | No. | Hydrops. | Discontinuation of treatment because of respiratory insufficiency and brain injury following asphyxia. |
| 34.0 | 0 | Yes. | Cystic kidney dysplasia, lung hypoplasia (anhydramnios), cardiomegaly, ascites, sandal gap. | Pulmonary hypoplasia |

| | | | | |
|---|---|---|---|---|
| 34.0 | 0 | Yes. | No autopsy. Babygram showed abnormalities suggestive of Jeune syndrome. | Palliative treatment. |
| 34.0 | 0 | No. | Omphalocele, bladder exstrophy, anorectal malformation, meningomyelocele, ventricular septal defect, hypoplastic lungs. | Palliative care because of multiple abnormalities. |
| 34.1 | 12 | No. | Hydrops, dysplastic kidneys, pyloric stenosis, uterus bicornis, dysmorphic facial features, rocker bottom feet. | Discontinuation of treatment because of multiple abnormalities. |
| 34.6 | 0 | Yes. | Laryngeal atresia, esophageal atresia, truncus arteriosus. | Unsuccessful resuscitation due to unexpected high airway obstruction. |
| 34.6 | 5 | No. | Hydrops, ventricular septal defect. | Sepsis and multiorgan dysfunction. |
| 34.7 | 0 | No. | No autopsy. Advanced anomaly scan: bilateral multicystic dysplastic kidneys and anhydramnios. | Palliative care because of infaust prognosis. |
| 35.1 | 0 | Yes. | Anorectal malformation, bilateral multicystic dysplastic kidneys, tracheoesophageal fistula. | Respiratory insufficiency due to pulmonary hypoplasia. |
| 36.3 | 4 | No. | Hydrocephalus, cerebellar dentate dysplasia, neuronal heterotopia. | Discontinuation of treatment because of hydrocephalus, severe pulmonary hypertension and convulsions. |
| 36.7 | 4 | Yes. | Hepatic haemangioendothelioma, cardiomegaly. | High-output failure. |
| 37.3 | 0 | No. | Coarctation of the aorta, atrial septal defect, mild pulmonary hypoplasia. | Respiratory insufficiency. |
| 37.7 | 0 | Yes. | None. | Multiorgan dysfunction after perinatal asphyxia. |
| 38.6 | 0 | No. | Abdominal neuroblastoma, multiple liver metastases, perimembranous ventricular septal defect. | Hypovolemic shock due to hemorrhage from abdominal tumor. |
| 38.9 | 0 | Yes. | Omphalocele, left diaphragmatic hernia, persistent left superior vena | Palliative care because of multiple abnormalities. |

| | | | | |
|---|---|---|---|---|
| | | | cava, absent processus xiphoideus, facial dysmorphic features, reduction defect left hand. | |
| 39.1 | 0 | No. | Ebstein’s anomaly | Cardiorespiratory failure. |
| 40.3 | 70 | No. | None. | Sudden infant death syndrome. |
| 40.9 | 1 | Yes. | None. | Subgaleal hemorrhage leading to cardiorespiratory failure. |
| 41.1 | 1 | Yes. | Small ventricular septal defect. | Sudden and unexpected postnatal collapse. |
| 41.3 | 25 | Yes. | Club feet, neuronal heterotopia. | Discontinuation of treatment due to poor prognosis. Neurological abnormalities (facial nerve paresis, bilateral vocal cord paresis, bilateral hearing loss). Suggestive of variation of Moebius syndrome. |

eci = e causa ignota, GA = Gestational age, IRDS = Infant respiratory distress syndrome, NEC = Necrotizing enterocolitis, PPROM = preterm premature rupture of membranes

**Table S3. The prevalence of cervical ribs and different vertebral patterns in fetuses and neonates with pathogenic chromosomal abnormalities, likely deleterious CNVs and variants of uncertain significance (N=374). One patient can have multiple CNVs, but is included only once in the most severe category.**

| Classification result chromosomal/ CNV analysis | Number of fetuses and neonates (% of total) | Cervical ribs (% within subgroup) | Vertebral pattern (number) |
|---|---|---|---|
| **Pathogenic** | | | |
| Aneuploidies | 15 (4.0) | 10 (66.7) | |
| Trisomy 21 | 7 (1.9) | 4 (57.1) | R (2), TL (1), CT (2), CT_TL (2) |
| Trisomy 13 | 3 (0.8) | 1 (33.3) | R (1), CT_TL_LS (2) |
| Trisomy 18 | 2 (0.5) | 2 (100) | CT_TL (1), CT_TL_LS (1) |
| Monosomy X | 2 (0.5) | 2 (100) | CT (1), CT_LS (1) |
| 47, XYY | 1 (0.3) | 1 (100) | CT (1) |
| Triploidy | 1 (0.3) | 0 | TL (1) |
| Pathogenic CNVs | 12 (3.2) | 6 (50.0%) | R (3)<br>TL (3)<br>CT (2)<br>CT_LS (1)<br>CT_TL (3) |
| **(Likely) deleterious** | | | |
| Large rare CNVs | 7/374 (1.9) | 4 (57.1) | R (2)<br>LS (1)<br>CT (1)<br>CT_TL (2)<br>CT_TL_LS (1) |
| **VOUS** | | | |
| Rare small CNVs | 37/374 (9.9) | 23 (62.2) | R (6)<br>LS (2)<br>TL (5)<br>TL_LS (1)<br>CT (7)<br>CT_LS (2)<br>CT_TL (12)<br>CT_TL_LS (2) |

CNVs = Copy number variants, VOUS = Variants of uncertain significance, R = Regular pattern, CT = shift at the cervicothoracic boundary, CT_TL = shift at the cervicothoracic and thoracolumbar boundary,

CT_LS = shift at cervicothoracic and lumbosacral boundary, TL_LS = shift at thoracolumbar and lumbosacral boundary, CT_TL_LS = shift at cervicothoracic, thoracolumbar and lumbosacral boundary.

**Table S4. Rare CNVs in the cohort and the dominant disease genes they have overlap with.**

| Patient | Candidate gene | Chromosome Region | Event | Length | Cytoband | Classification |
|---|---|---|---|---|---|---|
| 1 | - | chr18:18,539,806-18,785,222 | CN Loss | 245417 | q11.1 | VOUS |
| 2 | - | chr1:100,657,536-100,715,618 | CN Loss | 58083 | p21.2 | VOUS |
| 3 | - | chr17:0-737,451 | CN Loss | 737452 | p13.3 | VOUS |
| 4 | - | chr3:95,371,136-95,498,992 | CN Loss | 127857 | q11.2 | VOUS |
| 5 | - | chr17:1,919,027-2,105,149 | CN Loss | 186123 | p13.3 | VOUS |
| 6 | - | chr12:103,917,165-104,129,020 | CN Loss | 211856 | q23.3 | VOUS |
| 7 | - | chr17:50,398,465-50,882,343 | CN Gain | 483879 | q22 | Likely Benign |
| 8 | - | chr18:0-15,375,878 | CN Loss | 15375879 | p11.32 - p11.21 | Deleterious |
| 9 | - | chr7:23,340,039-23,500,438 | CN Gain | 160400 | p15.3 | VOUS |
| 10 | - | chr3:8,144,655-8,272,660 | CN Gain | 128006 | p26.1 | VOUS |
| 10 | FANCD2 | chr3:9,904,438-10,169,302 | CN Gain | 264865 | p25.3 | Likely deleterious |
| 11 | - | chr1:58,050,652-58,215,035 | CN Gain | 164384 | p32.2 | VOUS |
| 11 | - | chrX:710,944-1,326,153 | CN Gain | 615210 | p22.33 | VOUS |
| 12 | - | chr20:29,843,182-29,940,783 | CN Loss | 97602 | q11.21 | VOUS |
| 13 | - | chr22:21,037,716-21,630,630 | CN Loss | 592915 | q11.21 | Likely benign |
| 14 | - | chr11:20,559,294-20,733,949 | CN Loss | 174656 | p15.1 | VOUS |
| 15 | - | chr8:113,589,096-113,652,459 | CN Gain | 63364 | q23.3 | VOUS |
| 16 | - | chr9:73,961,653-74,019,112 | CN Gain | 57460 | q21.12 - q21.13 | Likely Benign |
| 17 | - | chr2:44,978,583-45,178,172 | CN Loss | 199590 | p21 | VOUS |
| 18 | GLI3 | chr7:42,104,194-42,215,177 | CN Loss | 110984 | p14.1 | VOUS |
| 19 | - | chr6:167,629,311-171,115,067 | CN Loss | 3485757 | q27 | Deleterious |
| 20 | - | chrX:21,367,202-21,462,735 | CN Gain | 95534 | p22.12 | VOUS |
| 21 | - | chr15:30,365,750-32,514,428 | CN Gain | 2148679 | q13.2 - q13.3 | VOUS |
| 22 | - | chr15:23,664,459-28,533,408 | CN Loss | 4868950 | q11.2 - q13.1 | Deleterious |
| 23 | - | chr5:75,072,512-75,203,710 | CN Loss | 131199 | q13.3 | Likely Benign |
| 24 | TBX1 | chr22:18,892,358-21,630,630 | CN Loss | 2738273 | q11.21 | Deleterious |
| 25 | - | chr1:15,873,540-15,936,834 | CN Loss | 63295 | p36.21 | VOUS |
| 25 | FANCA | chr16:87,999,977-89,545,160 | CN Loss | 1545184 | q24.2 - q24.3 | Likely deleterious |

| | | | | | | |
|---|---|---|---|---|---|---|
| 26 | - | chr7:100,936,290-101,185,980 | CN Gain | 249691 | q22.1 | Likely benign |
| 26 | - | chr22:47,738,154-47,891,583 | CN Loss | 153430 | q13.31 | Likely benign |
| 27 | - | chr10:120,943,898-121,131,977 | CN Loss | 188080 | q26.11 | VOUS |
| 28 | - | chr4:147,967,442-148,875,864 | CN Gain | 908423 | q31.22 - q31.23 | VOUS |
| 28 | - | chrX:1,722,313-2,127,376 | CN Gain | 405064 | p22.33 | VOUS |
| 29 | - | chrX:100,642,227-100,726,083 | CN Gain | 83857 | q22.1 | VOUS |
| 30 | - | chr15:86,405,874-87,092,508 | CN Gain | 686635 | q25.3 | VOUS |
| 31 | - | chr6:164,768,014-171,115,067 | CN Loss | 6347054 | q27 | Deleterious |
| 31 | - | chr9:0-22,938,918 | CN Gain | 22938919 | p24.3 - p21.3 | Deleterious |
| 32 | - | chr1:52,710,984-52,833,590 | CN Loss | 122607 | p32.3 | VOUS |
| 33 | - | chr7:155,894,750-157,022,646 | CN Loss | 1127897 | q36.3 | Likely deleterious |
| 34 | - | chr13:71,581,000-71,736,860 | CN Loss | 155861 | q21.33 | VOUS |
| 35 | - | chr5:179,350,039-180,915,260 | CN Loss | 1565222 | q35.3 | Likely deleterious |
| 35 | FANCA | chr16:87,803,969-90,354,753 | CN Gain | 2550785 | q24.2 - q24.3 | Likely deleterious |
| 36 | - | chr12:70,310,320-70,408,380 | CN Loss | 98061 | q15 | VOUS |
| 37 | - | chr1:90,535,793-90,660,638 | CN Gain | 124846 | p22.2 | Likely Benign |
| 38 | - | chr8:0-6,974,050 | CN Loss | 6974051 | p23.3 - p23.1 | Deleterious |
| 38 | - | chr8:11,856,723-43,646,413 | CN Gain | 31789690 | p23.1 - p11.22 | Deleterious |
| 39 | - | chr14:83,458,263-83,629,684 | CN Loss | 171422 | q31.1 - q31.2 | Likely Benign |
| 40 | - | chr21:34,860,460-34,977,465 | CN Loss | 117006 | q22.11 | VOUS |
| 41 | - | chr4:171,702,920-178,163,966 | CN Loss | 6461047 | q33 - q34.3 | Deleterious |
| 42 | - | chrX:42,980,588-43,061,007 | Homozygous Copy Loss | 80420 | p11.3 | VOUS |
| 43 | - | chr14:86,663,626-87,153,592 | CN Loss | 489967 | q31.3 | Likely Benign |
| 44 | - | chr16:65,256,521-65,310,477 | CN Loss | 53957 | q21 | Likely Benign |
| 45 | - | chr2:174,223,942-174,745,511 | CN Gain | 521570 | q31.1 | VOUS |
| 46 | - | chr2:31,472,334-31,542,953 | CN Loss | 70620 | p23.1 | VUS |
| 46 | - | chr9:94,233,244-94,325,779 | CN Gain | 92536 | q22.31 | Likely Benign |
| 47 | - | chr7:127,292,713-132,161,534 | CN Loss | 4868822 | q32.1 - q32.3 | Deleterious |
| 48 | - | chr1:10,735,816-10,805,288 | CN Gain | 69473 | p36.22 | VOUS |

| | | | | | | |
|---|---|---|---|---|---|---|
| 49 | - | chr8:145,966,164-146,193,436 | CN Gain | 227273 | q24.3 | VOUS |
| 50 | SHOX | chrX:0-2,673,491 | CN Gain | 2673492 | p22.33 | Likely deleterious |
| 51 | - | chr14:90,386,163-90,516,694 | CN Loss | 130532 | q32.11 | VOUS |
| 52 | - | chrX:1,687,084-2,183,615 | CN Gain | 496532 | p22.33 | VOUS |
| 53 | - | chr9:105,031,809-111,044,941 | CN Loss | 6013133 | q31.1 - q31.2 | Deleterious |
| 54 | - | chr2:26,576,134-26,776,189 | CN Gain | 200056 | p23.3 | VOUS |
| 54 | - | chr7:152,449,092-153,720,431 | CN Loss | 1271340 | q36.1 - q36.2 | Likely deleterious |
| 55 | - | chrX:6,450,700-8,138,035 | CN Loss | 1687336 | p22.31 | Likely deleterious |
| 56 | - | chr5:0-1,312,106 | CN Loss | 1312107 | p15.33 | Likely deleterious |
| 56 | - | chr10:0-4,905,883 | CN Gain | 4905884 | p15.3 - p15.1 | Deleterious |
| 57 | - | chr2:44,414,216-44,479,843 | CN Gain | 65628 | p21 | VOUS |
| 58 | - | chr22:34,740,652-34,807,627 | CN Loss | 66976 | q12.3 | Likely Benign |
| 59 | - | chr3:24,421,737-24,474,896 | CN Loss | 53160 | p24.2 | VOUS |
| 60 | - | chr6:170,544,940-171,115,067 | CN Loss | 570128 | q27 | Deleterious |
| 60 | - | chr13:80,439,088--115,169,878 | CN Gain | 34730790 | q31.1 -q34 | Deleterious |

CN = copy number, CNV = copy number variant, VOUS = Variants of uncertain significance

**Table S5. Genes that are involved in the rare CNVs identified in the cohort of deceased fetuses and neonates.**

| Name gene | chrom | strand |
|---|---|---|
| ABCA1 | 9 | - |
| ACER2 | 9 | + |
| ACSF3 | 16 | + |
| ACTR3B | 7 | + |
| ADAM18 | 8 | + |
| ADAM2 | 8 | - |
| ADAM28 | 8 | + |
| ADAM29 | 4 | + |
| ADAM32 | 8 | + |
| ADAM3A | 8 | - |
| ADAM5 | 8 | + |
| ADAM7 | 8 | + |
| ADAM9 | 8 | + |
| ADAMDEC1 | 8 | + |
| ADAMTSL1 | 9 | + |
| ADARB2 | 10 | - |
| ADARB2-AS1 | 10 | + |
| ADCYAP1 | 18 | + |
| ADGRA2 | 8 | + |
| ADRA1A | 8 | - |
| ADRB3 | 8 | - |
| AFDN | 6 | + |
| AFDN-AS1 | 6 | - |
| AFG3L1P | 16 | + |
| AFG3L2 | 18 | - |
| AGBL1 | 15 | + |
| AGBL1-AS1 | 15 | - |
| AGMAT | 1 | - |
| AGPAT5 | 8 | + |
| AHCYL2 | 7 | + |
| AHRR | 5 | + |
| AIFM3 | 22 | + |
| AK3 | 9 | - |
| AKAIN1 | 18 | - |
| AKAP17A | X | + |
| AKR1E2 | 10 | + |
| ANGPT2 | 8 | - |
| ANK1 | 8 | - |
| ANKRD11 | 16 | - |

| | | |
|---|---|---|
| ANKRD12 | 18 | + |
| ANKRD20A5P | 18 | + |
| ANKRD30B | 18 | + |
| ANKRD62 | 18 | + |
| AP3M2 | 8 | + |
| APCDD1 | 18 | + |
| APRT | 16 | - |
| ARHGAP10 | 4 | + |
| ARHGAP11B | 15 | + |
| ARHGAP28 | 18 | + |
| ARHGEF10 | 8 | + |
| ARMCX4 | X | + |
| ARVCF | 22 | - |
| ASAH1 | 8 | - |
| ASB5 | 4 | - |
| ASH2L | 8 | + |
| ASMT | X | + |
| ASMTL | X | - |
| ASMTL-AS1 | X | + |
| ATP10A | 15 | - |
| ATP6V1B2 | 8 | + |
| ATP6V1F | 7 | + |
| BAG4 | 8 | + |
| BANP | 16 | + |
| BCRP2 | 22 | + |
| BIN3 | 8 | - |
| BIN3-IT1 | 8 | - |
| BMP1 | 8 | + |
| BNC2 | 9 | - |
| BNIP3L | 8 | + |
| BRD9 | 5 | - |
| BRF2 | 8 | - |
| BRK1 | 3 | + |
| BTK | X | - |
| BTNL3 | 5 | + |
| BTNL8 | 5 | + |
| BTNL9 | 5 | + |
| C17orf97 | 17 | + |
| C18orf61 | 18 | + |
| C22orf29 | 22 | - |
| C22orf39 | 22 | - |
| C6orf118 | 6 | - |
| C6orf120 | 6 | + |

| | | |
|---|---|---|
| C7orf13 | 7 | - |
| C8orf4 | 8 | + |
| C8orf48 | 8 | + |
| C8orf58 | 8 | + |
| C8orf86 | 8 | - |
| C9orf66 | 9 | - |
| C9orf92 | 9 | - |
| CA5A | 16 | - |
| CALU | 7 | + |
| CAMKMT | 2 | + |
| CASZ1 | 1 | - |
| CBFA2T3 | 16 | - |
| CBWD1 | 9 | - |
| CBX3P2 | 18 | - |
| CC2D1B | 1 | - |
| CCAR2 | 8 | + |
| CCDC127 | 5 | - |
| CCDC136 | 7 | + |
| CCDC171 | 9 | + |
| CCDC188 | 22 | - |
| CCDC25 | 8 | - |
| CCR6 | 6 | + |
| CD274 | 9 | + |
| CD99 | X | + |
| CD99P1 | X | + |
| CDC37L1 | 9 | + |
| CDC37L1-AS1 | 9 | - |
| CDC45 | 22 | + |
| CDCA2 | 8 | + |
| CDCA7 | 2 | + |
| CDH15 | 16 | + |
| CDK10 | 16 | + |
| CDKN2A | 9 | - |
| CDKN2A-AS1 | 9 | + |
| CDKN2B | 9 | - |
| CDKN2B-AS1 | 9 | + |
| CDT1 | 16 | + |
| CENPBD1 | 16 | - |
| CEP192 | 18 | + |
| CEP41 | 7 | - |
| CEP44 | 4 | + |
| CEP72 | 5 | + |
| CEP76 | 18 | - |

| | | |
|---|---|---|
| CER1 | 9 | - |
| CETN1 | 18 | + |
| CHMP1A | 16 | - |
| CHMP1B | 18 | + |
| CHMP7 | 8 | + |
| CHRFAM7A | 15 | - |
| CHRNA2 | 8 | - |
| CHRNA6 | 8 | - |
| CHRNA7 | 15 | + |
| CHRNB3 | 8 | + |
| CIDEA | 18 | + |
| CIDEC | 3 | - |
| CIDECP | 3 | - |
| CLDN5 | 22 | - |
| CLN8 | 8 | + |
| CLTCL1 | 22 | - |
| CLU | 8 | - |
| CLUL1 | 18 | + |
| CNKSR2 | X | + |
| CNOT6 | 5 | + |
| CNOT7 | 8 | - |
| CNTLN | 9 | + |
| COL26A1 | 7 | + |
| COLEC12 | 18 | - |
| COMMD5 | 8 | - |
| COMT | 22 | + |
| COPG2 | 7 | - |
| CPA1 | 7 | + |
| CPA2 | 7 | + |
| CPA4 | 7 | + |
| CPA5 | 7 | + |
| CPNE7 | 16 | + |
| CRELD1 | 3 | + |
| CRKL | 22 | + |
| CRLF2 | X | - |
| CRYZL1 | 21 | - |
| CSF2RA | X | + |
| CSGALNACT1 | 8 | - |
| CSMD1 | 8 | - |
| CSMD3 | 8 | - |
| CTC-338M12.4 | 5 | + |
| CTD-3080P12.3 | 5 | - |
| CTU2 | 16 | + |

| | | |
|---|---|---|
| CXADRP3 | 18 | - |
| CYBA | 16 | - |
| CYLC2 | 9 | + |
| CYP4F35P | 18 | + |
| DAB1 | 1 | - |
| DACT2 | 6 | - |
| DBIL5P | 17 | + |
| DBNDD1 | 16 | - |
| DBT | 1 | - |
| DCTN6 | 8 | + |
| DDHD2 | 8 | + |
| DDX11L5 | 9 | + |
| DEF8 | 16 | + |
| DEFA1 | 8 | - |
| DEFA10P | 8 | - |
| DEFA11P | 8 | - |
| DEFA1B | 8 | - |
| DEFA3 | 8 | - |
| DEFA4 | 8 | - |
| DEFA5 | 8 | - |
| DEFA6 | 8 | - |
| DEFA8P | 8 | - |
| DEFA9P | 8 | - |
| DEFB1 | 8 | - |
| DEFB109P1 | 8 | - |
| DEFB115 | 20 | + |
| DEFB116 | 20 | - |
| DEFB130 | 8 | - |
| DEFB134 | 8 | - |
| DEFT1P | 8 | - |
| DEFT1P2 | 8 | - |
| DENND4C | 9 | + |
| DGCR10 | 22 | + |
| DGCR11 | 22 | - |
| DGCR14 | 22 | - |
| DGCR2 | 22 | - |
| DGCR5 | 22 | + |
| DGCR6 | 22 | + |
| DGCR6L | 22 | - |
| DGCR8 | 22 | + |
| DGCR9 | 22 | + |
| DHRSX | X | - |
| DIP2C | 10 | - |

| | | |
|---|---|---|
| DKFZP434L187 | 15 | + |
| DKK4 | 8 | - |
| DLC1 | 8 | - |
| DLGAP1 | 18 | - |
| DLGAP1-AS1 | 18 | + |
| DLGAP1-AS2 | 18 | + |
| DLGAP1-AS3 | 18 | + |
| DLGAP1-AS4 | 18 | + |
| DLGAP1-AS5 | 18 | + |
| DLGAP2 | 8 | + |
| DLGAP2-AS1 | 8 | - |
| DLL1 | 6 | - |
| DMRT1 | 9 | + |
| DMRT2 | 9 | + |
| DMRT3 | 9 | + |
| DMRTA1 | 9 | + |
| DMTN | 8 | + |
| DNAJC16 | 1 | + |
| DNAJC28 | 21 | - |
| DOC2B | 17 | - |
| DOCK5 | 8 | + |
| DOCK8 | 9 | + |
| DOK2 | 8 | - |
| DONSON | 21 | - |
| DPEP1 | 16 | + |
| DPH1 | 17 | + |
| DPP6 | 7 | + |
| DPYSL2 | 8 | + |
| DRC1 | 2 | + |
| DUSP26 | 8 | - |
| DUSP4 | 8 | - |
| EBF2 | 8 | - |
| EDNRA | 4 | + |
| EFCAB11 | 14 | - |
| EGR3 | 8 | - |
| EHD3 | 2 | + |
| EIF4EBP1 | 8 | + |
| ELP3 | 8 | + |
| EMC3 | 3 | - |
| EMC3-AS1 | 3 | + |
| EMILIN2 | 18 | + |
| ENOSF1 | 18 | - |
| ENTPD4 | 8 | - |

| | | |
|---|---|---|
| EPB41L3 | 18 | - |
| EPHX2 | 8 | + |
| ERICH1 | 8 | - |
| ERICH1-AS1 | 8 | + |
| ERLIN2 | 8 | + |
| ERMARD | 6 | + |
| ERMP1 | 9 | - |
| ESCO2 | 8 | + |
| EXOC3 | 5 | + |
| EXOC3-AS1 | 5 | - |
| EXTL3 | 8 | + |
| EXTL3-AS1 | 8 | - |
| FAM120B | 6 | + |
| FAM138C | 9 | - |
| FAM157C | 16 | + |
| FAM160B2 | 8 | + |
| FAM183CP | 8 | + |
| FAM210A | 18 | - |
| FAM230A | 22 | + |
| FAM230B | 22 | + |
| FAM57A | 17 | + |
| FAM66A | 8 | + |
| FAM66D | 8 | + |
| FAM71F1 | 7 | + |
| FAM71F2 | 7 | + |
| FAM86B1 | 8 | - |
| FAM86B2 | 8 | - |
| FAM87A | 8 | - |
| FAM90A25P | 8 | - |
| FAM90A2P | 8 | - |
| FAN1 | 15 | + |
| FANCA | 16 | - |
| FANCD2 | 3 | + |
| FANCD2OS | 3 | - |
| FBXO16 | 8 | - |
| FBXO25 | 8 | + |
| FBXO8 | 4 | - |
| FGF17 | 8 | + |
| FGF20 | 8 | - |
| FGFR1 | 8 | - |
| FGFR1OP | 6 | + |
| FGL1 | 8 | - |
| FKTN | 9 | + |

| | | |
|---|---|---|
| FLJ41200 | 9 | - |
| FLNC | 7 | + |
| FLT4 | 5 | - |
| FNTA | 8 | + |
| FOCAD | 9 | + |
| FOCAD-AS1 | 9 | - |
| FOXD4 | 9 | - |
| FREM1 | 9 | - |
| FRMD1 | 6 | - |
| FSD1L | 9 | + |
| FUT10 | 8 | - |
| FZD3 | 8 | + |
| GABRA5 | 15 | + |
| GABRB3 | 15 | - |
| GABRG3 | 15 | + |
| GABRG3-AS1 | 15 | - |
| GACAT2 | 18 | - |
| GALNS | 16 | - |
| GALNT7 | 4 | + |
| GALNTL6 | 4 | + |
| GAPLINC | 18 | + |
| GART | 21 | - |
| GAS8 | 16 | + |
| GAS8-AS1 | 16 | - |
| GEMIN4 | 17 | - |
| GFPT2 | 5 | - |
| GFRA2 | 8 | - |
| GINS4 | 8 | + |
| GLA | X | - |
| GLDC | 9 | - |
| GLI3 | 7 | - |
| GLIS3 | 9 | - |
| GLIS3-AS1 | 9 | + |
| GLOD4 | 17 | - |
| GLRA3 | 4 | - |
| GNAL | 18 | + |
| GNB1L | 22 | - |
| GNRH1 | 8 | - |
| GOLGA6L2 | 15 | - |
| GOLGA7 | 8 | + |
| GOLGA8H | 15 | + |
| GOLGA8J | 15 | + |
| GOLGA8R | 15 | - |

| | | |
|---|---|---|
| GOLGA8T | 15 | + |
| GOT1L1 | 8 | - |
| GP1BB | 22 | + |
| GPAT4 | 8 | + |
| GPM6A | 4 | - |
| GPR31 | 6 | - |
| GRK5 | 10 | + |
| GS1-24F4.2 | 8 | + |
| GSC2 | 22 | - |
| GSR | 8 | - |
| GTF2E2 | 8 | - |
| GTPBP4 | 10 | + |
| GTPBP6 | X | - |
| HACD4 | 9 | - |
| HAND2 | 4 | - |
| HAND2-AS1 | 4 | + |
| HAUS6 | 9 | - |
| HEIH | 5 | - |
| HERC2 | 15 | - |
| HERC2P10 | 15 | + |
| HGC6.3 | 6 | - |
| HGSNAT | 8 | + |
| HIC1 | 17 | + |
| HILPDA | 7 | + |
| HIRA | 22 | - |
| HMBOX1 | 8 | + |
| HMGB2 | 4 | - |
| HNRNPH2 | X | + |
| HOOK3 | 8 | + |
| HPGD | 4 | - |
| HR | 8 | - |
| HRAT5 | 5 | - |
| HTRA4 | 8 | + |
| IDI1 | 10 | - |
| IDI2 | 10 | - |
| IDI2-AS1 | 10 | + |
| IDO1 | 8 | + |
| IDO2 | 8 | + |
| IFNA1 | 9 | + |
| IFNA10 | 9 | - |
| IFNA13 | 9 | - |
| IFNA14 | 9 | - |
| IFNA16 | 9 | - |

| | | |
|---|---|---|
| IFNA17 | 9 | - |
| IFNA2 | 9 | - |
| IFNA21 | 9 | - |
| IFNA22P | 9 | - |
| IFNA4 | 9 | - |
| IFNA5 | 9 | - |
| IFNA6 | 9 | - |
| IFNA7 | 9 | - |
| IFNA8 | 9 | + |
| IFNB1 | 9 | - |
| IFNE | 9 | - |
| IFNW1 | 9 | - |
| IFT22 | 7 | - |
| IGF2BP3 | 7 | - |
| IKBKB | 8 | + |
| IL17C | 16 | + |
| IL17RC | 3 | + |
| IL17RE | 3 | + |
| IL33 | 9 | + |
| IL3RA | X | + |
| IMPA2 | 18 | + |
| IMPDH1 | 7 | - |
| INSL4 | 9 | + |
| INSL6 | 9 | - |
| INTS10 | 8 | + |
| INTS9 | 8 | - |
| IPW | 15 | + |
| IRF5 | 7 | + |
| JAGN1 | 3 | + |
| JAK2 | 9 | + |
| KANK1 | 9 | + |
| KAT6A | 8 | - |
| KBTBD11 | 8 | + |
| KBTBD11-OT1 | 8 | + |
| KCNU1 | 8 | + |
| KCNV2 | 9 | + |
| KCP | 7 | - |
| KCTD9 | 8 | - |
| KDM4C | 9 | + |
| KIAA1456 | 8 | + |
| KIAA2026 | 9 | - |
| KIF13B | 8 | - |
| KIF25 | 6 | + |

| | | |
|---|---|---|
| KIF25-AS1 | 6 | - |
| KLF13 | 15 | + |
| KLF14 | 7 | - |
| KLF4 | 9 | - |
| KLF6 | 10 | - |
| KLHDC10 | 7 | + |
| KLHL22 | 22 | - |
| KLHL9 | 9 | - |
| L3MBTL4 | 18 | - |
| L3MBTL4-AS1 | 18 | + |
| LAMA1 | 18 | - |
| LARP4B | 10 | - |
| LDLRAD4 | 18 | + |
| LDLRAD4-AS1 | 18 | - |
| LEP | 7 | + |
| LEPROTL1 | 8 | + |
| LETM2 | 8 | + |
| LGI3 | 8 | - |
| LINC00102 | X | - |
| LINC00106 | X | + |
| LINC00200 | 10 | + |
| LINC00242 | 6 | - |
| LINC00244 | 7 | + |
| LINC00304 | 16 | + |
| LINC00348 | 13 | + |
| LINC00470 | 18 | - |
| LINC00473 | 6 | - |
| LINC00513 | 7 | + |
| LINC00526 | 18 | - |
| LINC00574 | 6 | + |
| LINC00583 | 9 | + |
| LINC00587 | 9 | + |
| LINC00589 | 8 | - |
| LINC00602 | 6 | + |
| LINC00667 | 18 | + |
| LINC00668 | 18 | - |
| LINC00681 | 8 | + |
| LINC00685 | X | + |
| LINC00700 | 10 | - |
| LINC00701 | 10 | - |
| LINC00702 | 10 | - |
| LINC00703 | 10 | + |
| LINC00704 | 10 | - |

| | | |
|---|---|---|
| LINC00705 | 10 | + |
| LINC00847 | 5 | + |
| LINC00895 | 22 | - |
| LINC00896 | 22 | + |
| LINC00929 | 15 | + |
| LINC01000 | 7 | + |
| LINC01006 | 7 | - |
| LINC01230 | 9 | + |
| LINC01231 | 9 | + |
| LINC01239 | 9 | + |
| LINC01254 | 18 | - |
| LINC01255 | 18 | + |
| LINC01287 | 7 | - |
| LINC01288 | 8 | + |
| LINC01311 | 22 | + |
| LINC01387 | 18 | + |
| LINC01443 | 18 | + |
| LINC01444 | 18 | - |
| LINC01492 | 9 | - |
| LINC01505 | 9 | + |
| LINC01509 | 9 | - |
| LINC01558 | 6 | - |
| LINC01584 | 15 | - |
| LINC01605 | 8 | - |
| LINC01615 | 6 | - |
| LINC01624 | 6 | + |
| LINC01637 | 22 | + |
| LINC01644 | 22 | - |
| LINC01660 | 22 | - |
| LINC01882 | 18 | - |
| LINC01925 | 18 | - |
| LINC01962 | 5 | - |
| LINC02091 | 17 | - |
| LINC02099 | 8 | + |
| LINC02138 | 16 | + |
| LINC02153 | 8 | + |
| LINC-PINT | 7 | - |
| LMBR1 | 7 | - |
| LMCD1-AS1 | 3 | - |
| LOC100128714 | 15 | + |
| LOC100128993 | 8 | - |
| LOC100129697 | 16 | + |
| LOC100130705 | 7 | + |

| | | |
|---|---|---|
| LOC100130964 | 8 | + |
| LOC100132062 | 5 | + |
| LOC100133267 | 8 | - |
| LOC100192426 | 18 | - |
| LOC100287015 | 8 | - |
| LOC100287036 | 16 | + |
| LOC100288152 | 5 | + |
| LOC100288637 | 15 | + |
| LOC100289495 | 6 | + |
| LOC100289580 | 16 | + |
| LOC100506122 | 4 | + |
| LOC100506388 | 17 | + |
| LOC100506688 | 5 | - |
| LOC100506990 | 8 | + |
| LOC100507071 | 8 | - |
| LOC100507156 | 8 | + |
| LOC100996325 | 5 | - |
| LOC101927168 | 18 | - |
| LOC101927188 | 18 | + |
| LOC101927410 | 18 | + |
| LOC101927501 | X | + |
| LOC101927746 | 7 | + |
| LOC101927752 | 8 | - |
| LOC101927762 | 10 | + |
| LOC101927793 | 16 | + |
| LOC101927815 | 8 | - |
| LOC101927817 | 16 | + |
| LOC101927859 | 22 | + |
| LOC101927964 | 10 | + |
| LOC101928058 | 8 | - |
| LOC101928314 | 4 | - |
| LOC101928509 | 4 | - |
| LOC101928523 | 9 | + |
| LOC101928551 | 4 | + |
| LOC101928590 | 4 | + |
| LOC101928782 | 7 | + |
| LOC101928807 | 7 | + |
| LOC101928880 | 16 | + |
| LOC101929066 | 8 | + |
| LOC101929084 | 12 | + |
| LOC101929172 | 8 | + |
| LOC101929237 | 8 | + |
| LOC101929294 | 8 | - |

| | | |
|---|---|---|
| LOC101929297 | 6 | + |
| LOC101929315 | 8 | - |
| LOC101929420 | 6 | + |
| LOC101929460 | 6 | - |
| LOC101929470 | 8 | - |
| LOC101929504 | 6 | - |
| LOC101929523 | 6 | + |
| LOC101929550 | 8 | - |
| LOC101929622 | 8 | + |
| LOC101929897 | 8 | - |
| LOC101930370 | 4 | - |
| LOC102467222 | 8 | + |
| LOC102723376 | 10 | + |
| LOC102723701 | 8 | - |
| LOC102723729 | 8 | - |
| LOC102724357 | 6 | + |
| LOC102724467 | 16 | - |
| LOC102724511 | 6 | + |
| LOC102725022 | 15 | + |
| LOC102725080 | 8 | + |
| LOC104968399 | 18 | - |
| LOC105371414 | 16 | + |
| LOC105371430 | 17 | - |
| LOC105371485 | 17 | + |
| LOC105371998 | 18 | + |
| LOC105375504 | 7 | - |
| LOC105375972 | 9 | + |
| LOC105376194 | 9 | + |
| LOC105376351 | 10 | - |
| LOC105376360 | 10 | + |
| LOC105376365 | 10 | + |
| LOC105378123 | 6 | - |
| LOC105378127 | 6 | - |
| LOC105378137 | 6 | - |
| LOC105378146 | 6 | - |
| LOC105379393 | 8 | + |
| LOC154449 | 6 | - |
| LOC254896 | 8 | + |
| LOC283710 | 15 | - |
| LOC284865 | 22 | - |
| LOC285804 | 6 | - |
| LOC285889 | 7 | - |
| LOC286059 | 8 | + |

| | | |
|---|---|---|
| LOC286083 | 8 | - |
| LOC339059 | 16 | + |
| LOC339685 | 22 | + |
| LOC340357 | 8 | - |
| LOC340512 | 9 | - |
| LOC389641 | 8 | + |
| LOC389705 | 9 | + |
| LOC392196 | 8 | - |
| LOC400553 | 16 | - |
| LOC401052 | 3 | - |
| LOC401286 | 6 | - |
| LOC401442 | 8 | + |
| LOC407835 | 7 | + |
| LOC441052 | 4 | - |
| LOC441178 | 6 | - |
| LOC644669 | 18 | - |
| LOC649352 | 8 | - |
| LOC727896 | 18 | - |
| LOC728024 | 8 | - |
| LOC729732 | 8 | - |
| LONRF1 | 8 | - |
| LOXL2 | 8 | - |
| LPIN2 | 18 | - |
| LPL | 8 | + |
| LRRC14B | 5 | + |
| LRRC30 | 18 | + |
| LRRC4 | 7 | - |
| LRRC74B | 22 | + |
| LSM1 | 8 | - |
| LURAP1L | 9 | + |
| LURAP1L-AS1 | 9 | - |
| LZTR1 | 22 | + |
| LZTS1 | 8 | - |
| LZTS1-AS1 | 8 | + |
| MAGEL2 | 15 | - |
| MAK16 | 8 | + |
| MALSU1 | 7 | + |
| MAPK9 | 5 | - |
| MBOAT4 | 8 | - |
| MC1R | 16 | + |
| MC2R | 18 | - |
| MC5R | 18 | + |
| MCPH1 | 8 | + |

| | | |
|---|---|---|
| MCPH1-AS1 | 8 | - |
| MEAT6 | 6 | - |
| MED15 | 22 | + |
| MEST | 7 | + |
| MESTIT1 | 7 | - |
| METTL2B | 7 | + |
| METTL4 | 18 | - |
| MGAT1 | 5 | - |
| MGC27345 | 7 | - |
| MICU3 | 8 | + |
| MIR101-2 | 9 | + |
| MIR1286 | 22 | - |
| MIR129-1 | 7 | + |
| MIR1302-9 | 9 | + |
| MIR1306 | 22 | + |
| MIR132 | 17 | - |
| MIR182 | 7 | - |
| MIR183 | 7 | - |
| MIR185 | 22 | + |
| MIR1913 | 6 | - |
| MIR211 | 15 | - |
| MIR212 | 17 | - |
| MIR29A | 7 | - |
| MIR29B1 | 7 | - |
| MIR31 | 9 | - |
| MIR3148 | 8 | - |
| MIR3152 | 9 | + |
| MIR3156-2 | 18 | + |
| MIR31HG | 9 | - |
| MIR320A | 8 | - |
| MIR335 | 7 | + |
| MIR340 | 5 | - |
| MIR3618 | 22 | + |
| MIR3622A | 8 | + |
| MIR3622B | 8 | - |
| MIR3674 | 8 | + |
| MIR3690 | X | + |
| MIR383 | 8 | - |
| MIR3926-1 | 8 | - |
| MIR3926-2 | 8 | + |
| MIR3939 | 6 | - |
| MIR3976 | 18 | + |
| MIR3976HG | 18 | + |

| | | |
|---|---|---|
| MIR4276 | 4 | + |
| MIR4287 | 8 | - |
| MIR4288 | 8 | - |
| MIR4317 | 18 | - |
| MIR4456 | 5 | - |
| MIR4457 | 5 | - |
| MIR4469 | 8 | - |
| MIR4473 | 9 | - |
| MIR4474 | 9 | - |
| MIR4508 | 15 | - |
| MIR4526 | 18 | + |
| MIR4635 | 5 | - |
| MIR4638 | 5 | - |
| MIR4644 | 6 | + |
| MIR4659A | 8 | + |
| MIR4659B | 8 | - |
| MIR4665 | 9 | + |
| MIR4715 | 15 | - |
| MIR4722 | 16 | - |
| MIR4761 | 22 | + |
| MIR4767 | X | + |
| MIR4799 | 4 | + |
| MIR486-1 | 8 | - |
| MIR486-2 | 8 | + |
| MIR491 | 9 | + |
| MIR5189 | 16 | + |
| MIR5190 | 18 | + |
| MIR548AO | 8 | - |
| MIR548G | 4 | - |
| MIR548H4 | 8 | - |
| MIR548T | 4 | + |
| MIR548V | 8 | - |
| MIR5692A2 | 8 | + |
| MIR5699 | 10 | - |
| MIR593 | 7 | + |
| MIR596 | 8 | + |
| MIR6072 | 10 | - |
| MIR6078 | 10 | + |
| MIR6082 | 4 | + |
| MIR6089 | X | + |
| MIR649 | 22 | - |
| MIR6501 | 21 | + |
| MIR651 | X | + |

| | | |
|---|---|---|
| MIR6718 | 18 | + |
| MIR6775 | 16 | - |
| MIR6788 | 18 | - |
| MIR6816 | 22 | - |
| MIR6841 | 8 | - |
| MIR6842 | 8 | + |
| MIR6843 | 8 | - |
| MIR6850 | 8 | - |
| MIR6876 | 8 | + |
| MIR7153 | 18 | - |
| MIR7160 | 8 | + |
| MIR8055 | 8 | - |
| MIR8078 | 18 | - |
| MIR8081 | 9 | + |
| MIR8089 | 5 | - |
| MIR96 | 7 | - |
| MKLN1 | 7 | + |
| MKLN1-AS | 7 | - |
| MKRN3 | 15 | + |
| MLANA | 9 | + |
| MLLT3 | 9 | - |
| MNX1 | 7 | - |
| MNX1-AS1 | 7 | + |
| MPC1 | 6 | - |
| MPDZ | 9 | - |
| MPPE1 | 18 | - |
| MRM3 | 17 | + |
| MRPL40 | 22 | + |
| MSR1 | 8 | - |
| MTAP | 9 | + |
| MTCL1 | 18 | + |
| MTHFD2P1 | 3 | - |
| MTMR10 | 15 | - |
| MTMR7 | 8 | - |
| MTUS1 | 8 | - |
| MVD | 16 | - |
| MYL12A | 18 | + |
| MYL12B | 18 | + |
| MYOM1 | 18 | - |
| MYOM2 | 8 | + |
| MYRFL | 12 | + |
| NAPG | 18 | + |
| NAT1 | 8 | + |

| | | |
|---|---|---|
| NAT2 | 8 | + |
| NDC80 | 18 | + |
| NDN | 15 | - |
| NDUFV2 | 18 | + |
| NDUFV2-AS1 | 18 | - |
| NEFL | 8 | - |
| NEFM | 8 | + |
| NELL1 | 11 | + |
| NFIB | 9 | - |
| NIPSNAP3A | 9 | + |
| NIPSNAP3B | 9 | + |
| NKD2 | 5 | + |
| NKX2-6 | 8 | - |
| NKX3-1 | 8 | - |
| NKX6-3 | 8 | - |
| NOM1 | 7 | + |
| NPAP1 | 15 | + |
| NPM2 | 8 | + |
| NRF1 | 7 | + |
| NRG1 | 8 | + |
| NRG1-IT1 | 8 | + |
| NRG1-IT3 | 8 | + |
| NSD3 | 8 | - |
| NUDT18 | 8 | - |
| NUGGC | 8 | - |
| NXN | 17 | - |
| OCA2 | 15 | - |
| OPN1SW | 7 | - |
| OR13C2 | 9 | - |
| OR13C3 | 9 | - |
| OR13C4 | 9 | - |
| OR13C5 | 9 | - |
| OR13C8 | 9 | + |
| OR13C9 | 9 | - |
| OR13D1 | 9 | + |
| OR13F1 | 9 | + |
| OR2V1 | 5 | - |
| OR2V2 | 5 | + |
| OR2Y1 | 5 | - |
| OR4F21 | 8 | - |
| OR4F3 | 5 | + |
| OTOF | 2 | - |
| OTUD7A | 15 | - |

| | | |
|---|---|---|
| OVCA2 | 17 | + |
| P2RX6 | 22 | + |
| P2RX6P | 22 | - |
| P2RY8 | X | - |
| PABPN1L | 16 | - |
| PBK | 8 | - |
| PCM1 | 8 | + |
| PDCD1LG2 | 9 | + |
| PDCD2 | 6 | - |
| PDCD6 | 5 | + |
| PDE10A | 6 | - |
| PDGFRL | 8 | + |
| PDLIM2 | 8 | + |
| PEBP4 | 8 | - |
| PFKP | 10 | + |
| PGM5P3-AS1 | 9 | + |
| PHF10 | 6 | - |
| PHYHIP | 8 | - |
| PI4KA | 22 | - |
| PI4KAP1 | 22 | - |
| PIEZO1 | 16 | - |
| PIEZO2 | 18 | - |
| PITRM1 | 10 | - |
| PITRM1-AS1 | 10 | + |
| PIWIL2 | 8 | + |
| PLAT | 8 | - |
| PLCXD1 | X | + |
| PLEKHA2 | 8 | + |
| PLEKHG4B | 5 | + |
| PLGRKT | 9 | - |
| PLIN2 | 9 | - |
| PLPP5 | 8 | - |
| PLPP6 | 9 | + |
| PLXNA4 | 7 | - |
| PNMA2 | 8 | - |
| PNOC | 8 | + |
| PNPLA4 | X | - |
| PODXL | 7 | - |
| POLB | 8 | + |
| POLR3D | 8 | + |
| POM121L4P | 22 | + |
| POMK | 8 | + |
| POTEA | 8 | + |

| | | |
|---|---|---|
| POTEC | 18 | - |
| PP7080 | 5 | - |
| PPM1B | 2 | + |
| PPP2CB | 8 | - |
| PPP2R2A | 8 | + |
| PPP2R3B | X | - |
| PPP3CC | 8 | + |
| PPP4R1 | 18 | - |
| PPP4R1-AS1 | 18 | + |
| PRDM7 | 16 | - |
| PRELID3A | 18 | + |
| PRMT9 | 4 | - |
| PRODH | 22 | - |
| PROSC | 8 | + |
| PRR18 | 6 | - |
| PRR26 | 10 | + |
| PRRT3 | 3 | - |
| PRRT3-AS1 | 3 | + |
| PRRT4 | 7 | - |
| PSD3 | 8 | - |
| PSIP1 | 9 | - |
| PSMB1 | 6 | - |
| PSMG2 | 18 | + |
| PTK2B | 8 | + |
| PTPN2 | 18 | - |
| PTPRD | 9 | - |
| PTPRD-AS1 | 9 | + |
| PTPRD-AS2 | 9 | + |
| PTPRM | 18 | + |
| PUDP | X | - |
| PUM3 | 9 | - |
| PURG | 8 | - |
| PWAR1 | 15 | + |
| PWAR4 | 15 | + |
| PWAR5 | 15 | + |
| PWARSN | 15 | + |
| PWRN1 | 15 | + |
| PWRN2 | 15 | - |
| PWRN3 | 15 | + |
| R3HCC1 | 8 | + |
| RAB11FIP1 | 8 | - |
| RAB12 | 18 | + |
| RAB31 | 18 | + |

| | | |
|---|---|---|
| RACK1 | 5 | - |
| RAD23B | 9 | + |
| RALBP1 | 18 | + |
| RALGAPA1P1 | 9 | + |
| RANBP1 | 22 | + |
| RANBP6 | 9 | - |
| RASGEF1C | 5 | - |
| RBM28 | 7 | - |
| RBPMS | 8 | + |
| RBPMS-AS1 | 8 | - |
| RCL1 | 9 | + |
| REEP4 | 8 | - |
| RFLNB | 17 | - |
| RFX3 | 9 | - |
| RFX3-AS1 | 9 | + |
| RHOBTB2 | 8 | + |
| RIC1 | 9 | + |
| RIMBP3 | 22 | - |
| RLN1 | 9 | - |
| RLN2 | 9 | - |
| RNASET2 | 6 | - |
| RNF122 | 8 | - |
| RNF130 | 5 | - |
| RNF166 | 16 | - |
| RNF170 | 8 | - |
| RNF32 | 7 | + |
| RNF5P1 | 8 | - |
| RNMT | 18 | + |
| ROCK1 | 18 | - |
| ROCK1P1 | 18 | + |
| RPH3AL | 17 | - |
| RPL13 | 16 | + |
| RPL23AP53 | 8 | - |
| RPL36A | X | + |
| RPL36A-HNRNPH2 | X | + |
| RPL8 | 8 | - |
| RPS6 | 9 | - |
| RPS6KA2 | 6 | - |
| RPS6KA2-AS1 | 6 | + |
| RPS6KA2-IT1 | 6 | - |
| RRAGA | 9 | + |
| RTN4R | 22 | - |

| | | |
|---|---|---|
| RTN4RL1 | 17 | - |
| SAP30 | 4 | + |
| SARAF | 8 | - |
| SAXO1 | 9 | - |
| SCARA3 | 8 | + |
| SCARA5 | 8 | - |
| SCARF2 | 22 | - |
| SCARNA8 | 9 | - |
| SCGB3A1 | 5 | - |
| SCRG1 | 4 | - |
| SDAD1P1 | 8 | - |
| SDHA | 5 | + |
| SEH1L | 18 | + |
| SELENOI | 2 | + |
| SEPT5 | 22 | + |
| SEPT5-GP1BB | 22 | + |
| SERPIND1 | 22 | + |
| SFRP1 | 8 | - |
| SFT2D1 | 6 | - |
| SFTPC | 8 | + |
| SGCZ | 8 | - |
| SH2D4A | 8 | + |
| SH3GL2 | 9 | + |
| SHOX | X | + |
| SIX3 | 2 | + |
| SIX3-AS1 | 2 | - |
| SLC12A7 | 5 | - |
| SLC18A1 | 8 | - |
| SLC1A1 | 9 | + |
| SLC20A2 | 8 | - |
| SLC22A31 | 16 | - |
| SLC24A2 | 9 | - |
| SLC25A1 | 22 | - |
| SLC25A37 | 8 | + |
| SLC25A6 | X | - |
| SLC35G4 | 18 | + |
| SLC39A14 | 8 | + |
| SLC44A1 | 9 | + |
| SLC6A18 | 5 | + |
| SLC6A19 | 5 | + |
| SLC6A5 | 11 | + |
| SLC7A2 | 8 | + |
| SLC7A4 | 22 | - |

| | | |
|---|---|---|
| SLC7A5 | 16 | - |
| SLC9A3 | 5 | - |
| SMARCA2 | 9 | + |
| SMC2 | 9 | + |
| SMC2-AS1 | 9 | - |
| SMCHD1 | 18 | + |
| SMG6 | 17 | - |
| SMIM18 | 8 | + |
| SMIM19 | 8 | + |
| SMKR1 | 7 | + |
| SMO | 7 | + |
| SMOC2 | 6 | + |
| SNAI3 | 16 | - |
| SNAI3-AS1 | 16 | + |
| SNAP29 | 22 | + |
| SNAPC3 | 9 | + |
| SND1 | 7 | + |
| SND1-IT1 | 7 | + |
| SNORD107 | 15 | + |
| SNORD108 | 15 | + |
| SNORD109A | 15 | + |
| SNORD109B | 15 | + |
| SNORD115-1 | 15 | + |
| SNORD115-10 | 15 | + |
| SNORD115-11 | 15 | + |
| SNORD115-12 | 15 | + |
| SNORD115-13 | 15 | + |
| SNORD115-14 | 15 | + |
| SNORD115-15 | 15 | + |
| SNORD115-16 | 15 | + |
| SNORD115-17 | 15 | + |
| SNORD115-18 | 15 | + |
| SNORD115-19 | 15 | + |
| SNORD115-2 | 15 | + |
| SNORD115-20 | 15 | + |
| SNORD115-21 | 15 | + |
| SNORD115-22 | 15 | + |
| SNORD115-23 | 15 | + |
| SNORD115-24 | 15 | + |
| SNORD115-25 | 15 | + |
| SNORD115-26 | 15 | + |
| SNORD115-27 | 15 | + |
| SNORD115-28 | 15 | + |

| | | |
|---|---|---|
| SNORD115-29 | 15 | + |
| SNORD115-3 | 15 | + |
| SNORD115-30 | 15 | + |
| SNORD115-31 | 15 | + |
| SNORD115-32 | 15 | + |
| SNORD115-33 | 15 | + |
| SNORD115-34 | 15 | + |
| SNORD115-35 | 15 | + |
| SNORD115-36 | 15 | + |
| SNORD115-37 | 15 | + |
| SNORD115-38 | 15 | + |
| SNORD115-39 | 15 | + |
| SNORD115-4 | 15 | + |
| SNORD115-40 | 15 | + |
| SNORD115-41 | 15 | + |
| SNORD115-42 | 15 | + |
| SNORD115-43 | 15 | + |
| SNORD115-44 | 15 | + |
| SNORD115-45 | 15 | + |
| SNORD115-46 | 15 | + |
| SNORD115-47 | 15 | + |
| SNORD115-48 | 15 | + |
| SNORD115-5 | 15 | + |
| SNORD115-6 | 15 | + |
| SNORD115-7 | 15 | + |
| SNORD115-8 | 15 | + |
| SNORD115-9 | 15 | + |
| SNORD116-1 | 15 | + |
| SNORD116-10 | 15 | + |
| SNORD116-11 | 15 | + |
| SNORD116-12 | 15 | + |
| SNORD116-13 | 15 | + |
| SNORD116-14 | 15 | + |
| SNORD116-15 | 15 | + |
| SNORD116-16 | 15 | + |
| SNORD116-17 | 15 | + |
| SNORD116-18 | 15 | + |
| SNORD116-19 | 15 | + |
| SNORD116-2 | 15 | + |
| SNORD116-20 | 15 | + |
| SNORD116-21 | 15 | + |
| SNORD116-22 | 15 | + |
| SNORD116-23 | 15 | + |

| | | |
|---|---|---|
| SNORD116-24 | 15 | + |
| SNORD116-25 | 15 | + |
| SNORD116-26 | 15 | + |
| SNORD116-27 | 15 | + |
| SNORD116-28 | 15 | + |
| SNORD116-29 | 15 | + |
| SNORD116-3 | 15 | + |
| SNORD116-30 | 15 | + |
| SNORD116-4 | 15 | + |
| SNORD116-5 | 15 | + |
| SNORD116-6 | 15 | + |
| SNORD116-7 | 15 | + |
| SNORD116-8 | 15 | + |
| SNORD116-9 | 15 | + |
| SNORD13 | 8 | + |
| SNORD137 | 9 | - |
| SNORD142 | 10 | + |
| SNORD64 | 15 | + |
| SNORD68 | 16 | + |
| SNORD95 | 5 | - |
| SNORD96A | 5 | - |
| SNRPN | 15 | + |
| SNURF | 15 | + |
| SON | 21 | + |
| SORBS3 | 8 | + |
| SPATA2L | 16 | - |
| SPATA33 | 16 | + |
| SPATA4 | 4 | - |
| SPATA6L | 9 | - |
| SPCS3 | 4 | + |
| SPG7 | 16 | + |
| SPIRE1 | 18 | - |
| SPIRE2 | 16 | + |
| SSMEM1 | 7 | + |
| STAB2 | 12 | + |
| STAR | 8 | - |
| STC1 | 8 | - |
| STMN4 | 8 | - |
| STRIP2 | 7 | + |
| STS | X | + |
| T | 6 | - |
| TACC1 | 8 | + |
| TAL2 | 9 | + |

| | | |
|---|---|---|
| TANGO2 | 22 | + |
| TBP | 6 | + |
| TBX1 | 22 | + |
| TCF25 | 16 | + |
| TCP10 | 6 | - |
| TCP10L2 | 6 | + |
| TCTE3 | 6 | - |
| TDP1 | 14 | + |
| TDRP | 8 | - |
| TERT | 5 | - |
| TEX15 | 8 | - |
| TGIF1 | 18 | + |
| THAP1 | 8 | - |
| THAP7 | 22 | - |
| THAP7-AS1 | 22 | + |
| THBS2 | 6 | - |
| THOC1 | 18 | - |
| THRB | 3 | - |
| TM2D2 | 8 | - |
| TMEM184C | 4 | + |
| TMEM191A | 22 | + |
| TMEM191B | 22 | + |
| TMEM200C | 18 | - |
| TMEM209 | 7 | - |
| TMEM261 | 9 | - |
| TMEM38B | 9 | + |
| TNFRSF10A | 8 | - |
| TNFRSF10B | 8 | - |
| TNFRSF10C | 8 | + |
| TNFRSF10D | 8 | - |
| TNPO3 | 7 | - |
| TPD52L3 | 9 | + |
| TPI1P2 | 7 | + |
| TPPP | 5 | - |
| TRAPPC2L | 16 | + |
| TRIM35 | 8 | - |
| TRIM41 | 5 | + |
| TRIM52 | 5 | - |
| TRIM52-AS1 | 5 | + |
| TRIM7 | 5 | - |
| TRIP13 | 5 | + |
| TRMT2A | 22 | - |
| TRPM1 | 15 | - |

| | | |
|---|---|---|
| TSGA13 | 7 | - |
| TSPAN33 | 7 | + |
| TSSK2 | 22 | + |
| TTC39B | 9 | - |
| TTI2 | 8 | - |
| TTLL2 | 6 | + |
| TUBA3FP | 22 | - |
| TUBB3 | 16 | + |
| TUBB6 | 18 | + |
| TUBB8 | 10 | - |
| TUSC3 | 8 | + |
| TWSG1 | 18 | + |
| TXNDC2 | 18 | + |
| TXNRD2 | 22 | - |
| TYMS | 18 | + |
| TYMSOS | 18 | - |
| TYRP1 | 9 | + |
| UBE2H | 7 | - |
| UBE3A | 15 | - |
| UBE3C | 7 | + |
| UBXN8 | 8 | + |
| UFD1L | 22 | - |
| UHRF2 | 9 | + |
| ULK4P2 | 15 | + |
| ULK4P3 | 15 | + |
| UNC5D | 8 | + |
| UNC93A | 6 | + |
| URAHP | 16 | - |
| USP14 | 18 | + |
| USP17L2 | 8 | - |
| USP17L7 | 8 | - |
| VAPA | 18 | + |
| VCX | X | + |
| VCX2 | X | - |
| VCX3A | X | - |
| VDAC3 | 8 | + |
| VEGFC | 4 | - |
| VLDLR | 9 | + |
| VLDLR-AS1 | 9 | - |
| VPS37A | 8 | + |
| VPS53 | 17 | - |
| VPS9D1 | 16 | - |
| VPS9D1-AS1 | 16 | + |

| | | |
|---|---|---|
| WASHC1 | 9 | - |
| WDR17 | 4 | + |
| WDR27 | 6 | - |
| WDR37 | 10 | + |
| WRN | 8 | + |
| XG | X | + |
| XKR5 | 8 | - |
| XPO7 | 8 | + |
| YES1 | 18 | - |
| ZBED1 | X | - |
| ZBTB14 | 18 | - |
| ZC3H18 | 16 | + |
| ZC3HC1 | 7 | - |
| ZDHHC11 | 5 | - |
| ZDHHC2 | 8 | + |
| ZDHHC21 | 9 | - |
| ZDHHC8 | 22 | + |
| ZFP62 | 5 | - |
| ZFPM1 | 16 | + |
| ZFYVE9 | 1 | + |
| ZMAT4 | 8 | - |
| ZMYND11 | 10 | + |
| ZNF16 | 8 | - |
| ZNF250 | 8 | - |
| ZNF251 | 8 | - |
| ZNF276 | 16 | + |
| ZNF34 | 8 | - |
| ZNF395 | 8 | - |
| ZNF462 | 9 | + |
| ZNF469 | 16 | + |
| ZNF517 | 8 | + |
| ZNF519 | 18 | - |
| ZNF596 | 8 | + |
| ZNF7 | 8 | + |
| ZNF703 | 8 | + |
| ZNF705D | 8 | + |
| ZNF74 | 22 | + |
| ZNF778 | 16 | + |

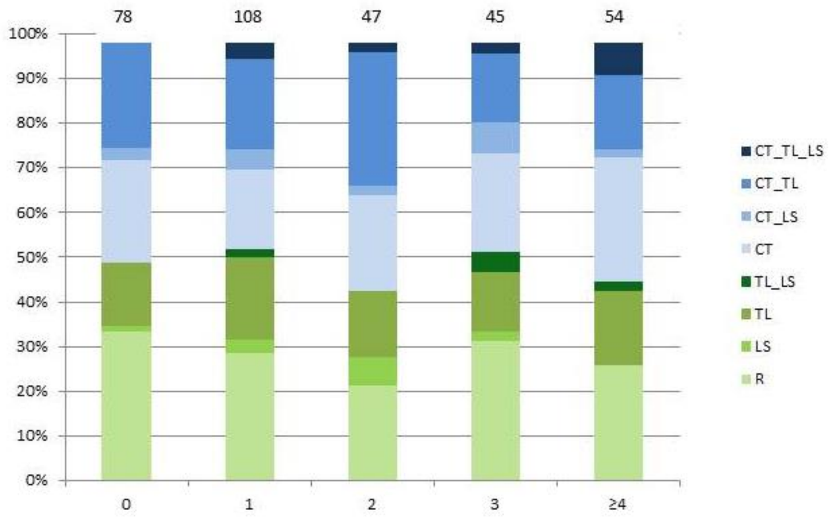


**Figure S1**. The pattern of the vertebral column in fetuses and neonates categorized according to the number of affected organ systems depicted as 0 (no anomalies) or 1, 2, 3 or ≥ 4 (number of affected organs sytems on the X bar). The total number of cases in each group is depicted above the bars.

R= Regular pattern, CT= shift at the cervicothoracic boundary, CT_TL= shift at the cervicothoracic and thoracolumbar boundary, CT_LS= shift at cervicothoracic and lumbosacral boundary, TL_LS= shift at thoracolumbar and lumbosacral boundary, CT_TL_LS= shift at cervicothoracic, thoracolumbar and lumbosacral boundary.

### Supplementary methods

*Analysis of copy number variation*

DNA was isolated from material that was collected in patients opting for invasive prenatal or postnatal diagnostic tests (e.g. amniocentesis, chorion villus biopsy, skin biopsy). Subsequently, these DNA samples were used to determine CNV profiles using several types of SNP array. (12-HumanCyto SNP DNA Analysis BeadChips v2.1, HumanOmniExpress BeadChip , Infinium CytoSNP-850k (Illumina Inc., San Diego, CA, USA ) and the GeneChip Human Mapping 250K Nsp (Affymetrix Inc. Santa Clara, CA, USA)). We determined the Copy Number variation (CNV) profiles in coding and non-coding regions of patients (n=265) using methods and analysis settings previously described.[1] CNV profiles were inspected visually in Biodiscovery Nexus

CN8.0. (Biodiscovery Inc., Hawthorne, CA, USA). Losses and gains were considered for evaluation when larger than 50kb and absent from large (n=19.585) control cohorts.[2, 3] Homozygous losses had to contain at least 6 probes. Rare coding CNV were inspected for gene content, rare noncoding CNV were inspected for putatively disrupting topologically associated domain (TAD) borders [4] or to contain enhancers.[5, 6] We classified the CNVs with an overlap of at least 75% with similar state CN changes (loss/gain) in the control cohort as likely benign. CNV absent from these cohorts as rare CNV. We used this stringent cut-off as cervical ribs are extremely rare and we expect contributing genetic defects to be absent in the normal population. Remaining rare CNVs were classified as described in table 1.

References


1. Brosens, E., et al., *Copy number variations in 375 patients with oesophageal atresia and/or tracheoesophageal fistula.* Eur J Hum Genet, 2016. **24**(12): p. 1715-1723.
2. Coe, B.P., et al., *Refining analyses of copy number variation identifies specific genes associated with developmental delay.* Nat Genet, 2014. **46**(10): p. 1063-71.
3. Cooper, G.M., et al., *A copy number variation morbidity map of developmental delay.* Nat Genet, 2011. **43**(9): p. 838-46.
4. Rao, S.S., et al., *A 3D map of the human genome at kilobase resolution reveals principles of chromatin looping.* Cell, 2014. **159**(7): p. 1665-80.
5. Visel, A., et al., *VISTA Enhancer Browser--a database of tissue-specific human enhancers.* Nucleic Acids Res, 2007. **35**(Database issue): p. D88-92.
6. Pennacchio, L.A., et al., *In vivo enhancer analysis of human conserved non-coding sequences.* Nature, 2006. **444**(7118): p. 499-502.
